# Oxytocin administration in neonates shapes the hippocampal circuitry and restores social behavior in a mouse model of autism

**DOI:** 10.1101/2020.09.21.306217

**Authors:** Alessandra Bertoni, Fabienne Schaller, Roman Tyzio, Stephane Gaillard, Francesca Santini, Marion Xolin, Diabé Diabira, Radhika Vaidyanathan, Valery Matarazzo, Igor Medina, Elizabeth Hammock, Jinwei Zhang, Bice Chini, Jean-Luc Gaiarsa, Françoise Muscatelli

**Affiliations:** Institute of Neurobiology of Méditerranée (INMED), Institut National de la Santé et de la Recherche Médicale (INSERM) UMR 1249, Aix- Marseille Université, Marseille, France; Phenotype-expertise, Marseille, France; Florida State University, Tallahassee, FL, USA; Institute of Neuroscience, National Research Council (CNR), Department of Medical Biotechnology and Translational Medicine, Università degli Studi di Milano Milan, and NeuroMI Milan Center for Neuroscience, University of Milano-Bicocca, Italy; Institute of Biomedical and Clinical Sciences, College of Medicine and Health, University of Exeter, Hatherly Laboratories, Exeter, EX4 4PS, UK

**Keywords:** Neurodevelopmental disorder, behavior, Excitation/Inhibition balance, genetics, MAGEL2, Schaaf-Yang and Prader-Willi Syndromes.

## Abstract

Oxytocin is a master regulator of the social brain. In some animal models of autism, notably in *Magel2 ^tm1.1Mus^*-deficient mice, peripheral administration of oxytocin in infancy improves social behaviors until adulthood. However, neither the mechanisms responsible for social deficits nor the mechanisms by which such oxytocin administration has long-term effects are known. Here, we aimed to clarify these oxytocin-dependent mechanisms focusing on social memory performance.

We showed that *Magel2^tm1.1Mus^-*deficient mice present a deficit in social memory and studied the hippocampal circuits underlying this memory. We showed a co-expression of *Magel2* and *oxytocin-receptor* in the dentate gyrus and CA2/CA3 hippocampal regions. Then, we demonstrated: an increase of the GABAergic activity of CA3-pyramidal cells associated with an increase in the quantity of oxytocin-receptors and of somatostatin interneurons. We also revealed a delay in the GABAergic development sequence in *Magel2^tm1.1Mus^*-deficient pups, linked to phosphorylation modifications of KCC2. Above all, we demonstrated the positive effects of subcutaneous administration of oxytocin in the mutant neonates, restoring neuronal alterations and social memory.

Although clinical trials are debated, this study highlights the mechanisms by which peripheral oxytocin-administration in neonates impacts the brain and demonstrates the therapeutic value of oxytocin to treat infants with autism spectrum disorders.

## INTRODUCTION

The nonapeptide oxytocin (OT) and its signaling pathway, the OT-system, is a master regulator in the development of the social brain suggesting that OT plays a role in both childhood and adult neuropsychiatric disorders characterized by social cognition impairment (1). OT-system is disrupted in several animal models of neurodevelopmental disorders (2, 3). Indeed, knockout mouse models of *oxytocin* (4, 5), *oxytocin-receptor* (*Oxtr*) (6–8), or *ADP-ribosyl cyclase* (*Cd38*) (9, 10) genes show changes in social behavior reminiscent of autism spectrum disorders (ASD). On the other way, several rodent models of ASD due either to the inactivation of genes such as *Fmr1, Cntnap2, Magel2, Oprm1, Shank3*, *Nlgn-3* or to environmental valproic acid exposure (VPA), exhibit indirectly a deficit of the brain OT-system (2). *MAGEL2* is a gene that is involved in Prader-Willi (PWS) (11) and Schaaf-Yang (SYS) syndromes (12) and is classified as one of the highest relevant gene to ASD risk (SFARI ranking). Both of these genetic neurodevelopmental disorders have in common autistic features with alterations in social behavior and deficits in cognition that persist over the lifespan (13). *Magel2^tm1.1Mus^-* deficient mouse model is a pertinent model for both syndromes (13), mimicking alterations in social behavior and learning abilities in adulthood (14, 15). *Magel2* is expressed in the developing hypothalamus until adulthood and *Magel2^tm1.1Mus^-* knockout (KO) neonates display a deficiency of several hypothalamic neuropeptides, particularly OT (16). Daily administration of OT in *Magel2-KO* neonates during the first week of life improves social behavior and learning abilities beyond treatment into adulthood (14). Comparable long term effects have also been reported in other genetic rodent models such as the VPA-induced rat model (17), the *Cntnap2* and *Fmr1* KO mice (18, 19), and following maternal separation (20). However, the neurobiological alterations involving the OT-system and responsible for social behavior deficits in these models are not known. Similarly, the mechanisms by which OT-treatment in infancy exerts its long-lasting beneficial effects, remain mysterious.

At adulthood, OT is thought to regulate aspects of social behavior via interactions with OXTRs in a number of key brain regions (21). Social recognition memory deficit in adulthood is the most robust phenotype, present in all the models with an alteration of the OT-system (22). With regard to social memory, a critical role has been ascribed to hippocampal OXTR expression in the anterior dentate gyrus (aDG) hilar and anterior CA2/CA3distal regions (aCA2/CA3d) (23–26). In the aCA2/CA3d region, OXTRs are expressed in glutamatergic pyramidal neurons and in GABAergic interneurons, which account for over 90% of OXTR positive cells in the hippocampus (27). Notably, both types of neuron are necessary for the formation of stronger synapses that mediate long term potentiation and social memory (23, 24, 28).

In the first two postnatal weeks, OT neuron projections set up and the expression of OXTRs is extremely dynamic followed by a decreased expression thereafter (29, 30). However, during this developmental period, the mechanisms by which the OT-system structures various behaviors are little studied. One study reports that OT is involved dendritic and synaptic refinement in immature hippocampal glutamatergic neurons (31).

Five clinical trials (phase 1 or 2) of OT administration in patients with PWS have been conducted and positive or no effects have been reported but no adverse effects (32). However, each of the studies is fairly empirical and uses different timings, durations and doses of OT, since we do not yet understand clearly how OT works. Based on our previous preclinical studies (14, 16), a phase 1/2 clinical trial with OT-treatment of infants with PWS significantly improves early feeding and “social skills” (33), supporting the translational relevance of our study. More research is needed to demonstrate and validate our hypothesis that the administration of OT in early infancy might be the most beneficial treatment for PWS/SYS. Thus, to build a strong scientific rationale, it is necessary to elucidate the PWS/SYS neuronal alterations and the mechanisms underlying the long-lasting effects of OT-administration in neonates.

Here, we aimed to clarify the physiological and cellular mechanisms related to the OT-system that are disturbed in *Magel2^tm1.1Mus^-deficient* mice and those responsible for the long-term rescue effects following OT peripheral administration in pups. We focused our study on the deficit of social memory, a robust phenotype linked to OT-system.

## METHODS AND MATERIALS

### Animals and primary hippocampal cultures

*Magel2^tm1.1Mus^*+/+ (WT) and *Magel2^tm1.1Mus^*-/- (*Magel2-KO*) mice were maintained on a C57BL/6J genetic background. Experimental protocols were approved by the institutional Ethical Committee guidelines for animal research with the accreditation no. B13-055-19 from the French Ministry of Agriculture. *Magel2*-deficient mice were generated as previously published (16). Embryonic day 18 dissociated hippocampal neurons were obtained from timed pregnant mice as previously described (34). See *Supplemental Information* for details.

### Oxytocin Treatment

WT and *Magel2-KO* pups were removed from their mother, placed on a heating pad, given a subcutaneous (s.c.) injection and quickly returned to the mother. The solutions injected were isotonic saline (10 µl) for control mice and 2 µg of OT (Phoenix Pharmaceuticals Inc., cat #051-01) diluted in isotonic saline (10 µl) for treated mice. The treatment was performed during the first week of life every other day (P0, P2, P4, P6).

### Behavior

The effects of *Magel2* deletion and OT-treatment were evaluated on social behavior, locomotor and vertical activity, anxiety and non-social memory. For detailed procedures, see *Supplemental Information*.

### Calcium imaging recordings

Calcium imaging experiments were carried out as previously reported (35) and are briefly described in *Supplemental Information* section.

### Electrophysiological recordings and morphological analysis

*Whole-cell patch clamp:* spontaneous and miniature synaptic activity was recorded in voltage-clamp mode on P20-P25 CA3 pyramidal neurons. The morphology of recorded aCA3d neurons was defined by adding biocytin in the recording solution and performing Neurolucida reconstruction followed by a Sholl analysis. See *Supplemental Information*, for details.

*Single GABA_A_ channel recordings* were performed on hippocampal CA3 pyramidal neurons at P1, P7 and P15 in cell-attached configuration, as described in *Supplemental Information*.

### Immunohistochemistry and quantification

Immunostaining was carried out on 50μm-thick coronal sections following standard procedure, as described in *Supplementary Information*.

### OT binding assay

Adult WT and mutant mice were sacrificed and non-perfused mouse brain were frozen in - 25°C isopentane and stored at −80°C until cut. 14 µm-thick brain slices were cut using a cryostat (Frigocut-2700, Reichert-Jung) and collected on chromallume-coated slides and stored at −80°C until use. Localization of OT binding sites was performed by autoradiography as previously described (36) and detailed in *Supplemental Information*.

### Chromogenic In situ Hybridization

Fresh-frozen brains from WT mice at P7 and P28 were sectioned in a cryostat in the coronal plane at 20μm thickness and mounted on Superfrost Plus slides and stored at −80°C. RNA detection was performed on tissue sections using RNAscope 2.5HD Duplex Assay (Cat #322430, Advanced Cell Diagnostics (ACD), Hayward, CA) as detailed in *Supplemental Information*.

### Western Blot

Western blotting experiments were performed on hippocampal tissue and specific bands were visualized with secondary HRP-conjugated antibodies using ChemiDoc™ Imaging Systems (Bio-Rad). The relative intensities of immunoblot bands were determined by densitometry with ImageJ software. See *Supplemental Information* for details.

### Statistical Analysis

Statistical analyses were performed using GraphPad Prism (GraphPad Software, Prism 7.0 software, Inc, La Jolla, CA, USA). All the statistical analyses are reported in a specific file. For details, see *Supplemental Information*.

## RESULTS

In the following experiments WT and *Magel2^tm1.1Mus^-*KO (*Magel2-*KO) pups were naïve or treated for four days (Postnatal day P0, P2, P4 and P6) with one subcutaneous administration of physiological saline (“vehicle”) or oxytocin (2 µg; “OT-treatment” or “+OT”) per day.

### Deficit of social memory in *Magel2-KO* males is rescued by neonatal OT-treatment

At adulthood, we focused on social behavior using the three-chamber test in order to assess social exploration (sociability), the preference for social novelty (social discrimination) and social memory (short-term social memory) (Figure 1A). *Magel2-*KO males showed levels of sociability and social discrimination similar to WT males but exhibited a significant deficit in social memory (Figure 1B, Figure 1-Supplement1). As reported (37), we observed a failure of the three-chamber test in revealing sociability in the cohort of female mice (Figure 1-Supplement2). As a consequence, we restricted all following studies to male mice.

**Figure 1:**
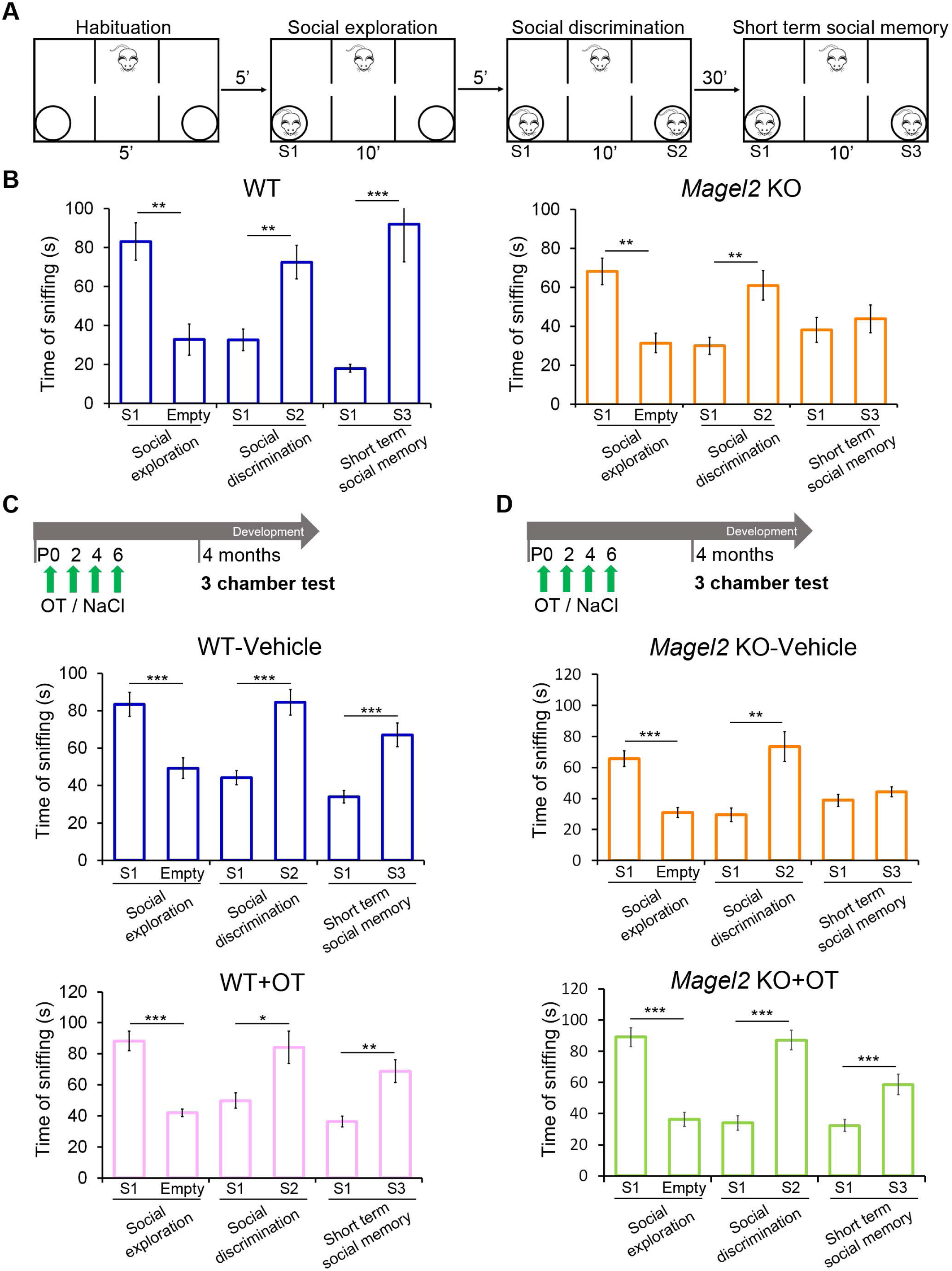
Social behavior in three-chamber test of male *Magel2* KO adult versus WT adults and having been OT-treated or vehicle-treated in neonates. **(A)** Paradigm of the three-chamber test. Sniffing time between mice is measured in each test. **(B)** WT males (N=9) show normal behavior in all the steps of the test; *Magel2* KO males (N=9) show a significant impairment in short term social memory. (**C**) WT mice were treated in the first week of life with vehicle or OT and then tested at four months. WT mice treated with vehicle (N=18) or treated with OT (N=10) have similar profiles with significant differences in each step of the test. (**D**) *Magel2* KO mice were treated in the first week of life with vehicle or OT and then tested at four months. *Magel2* KO-vehicle (N=19) mice show a significant difference in the social exploration and social discrimination, but, in short term social memory, they do not show a higher sniffing time with the novel mouse. OT-treated *Magel2* KO (N=19) mice present significant differences in each step of the step. Data represented in histograms report the interaction time (time of sniffing in seconds) as mean ± SEM. Mann-WhitneyTest. *P<0.05, **P<0.01, ***P<0.001. Statistical analysis is reported in Supplemental Table 1.

First, the effects of neonatal vehicle and OT-treatment were assessed in WT male pups at adulthood in the three-chamber test. We found that neither treatment had any measurable effect on sociability, social discrimination or social memory: the amount of time spent sniffing in different compartments was similar to that recorded in untreated WT males (Figures 1B-C). Furthermore, no significant effects of neonatal OT-treatment in WT animals were detected in widely used assays to test object recognition and social behavior (Figure1-Supplement3A,B), motor activity (Figure1-Supplement3C) and anxiety-like behaviors (Figure 1-Supplement3B,C). Then, neonatal vehicle and OT-treatment were administered in *Magel2*-KO pups. Unsurprisingly, *Magel2-*KO-vehicle males presented a social memory deficit similar to untreated *Magel2-*KO males (Figure 1B,D). However, this deficit was rescued by neonatal OT-treatment (Figure 1D). Sociability and social discrimination indices were not affected by vehicle or OT-treatment (Figure 1-Supplement1).

Thus, the loss of *Magel2* causes a deficit in social memory in male *Magel2-*KO adults. This deficit was rescued by a neonatal OT-treatment. Due to the robust effect observed on social memory, we focused our subsequent investigations on the hippocampal region, previously shown to be specifically involved in OT-mediated effects on social memory (23, 24). Neurons expressing the OXTRs in the CA2/CA3d and DG regions of the anterior hippocampus are involved in social memory (23, 24) therefore we first tested whether *Magel2* and *Oxtr* are co-expressed in those regions.

### aDG and aCA2/CA3d regions co-express *Magel2* and *Oxtr* transcripts

*Magel2* is known to be highly expressed in hypothalamus, while its expression in hippocampal regions is less well characterized. Taking in account the developmental and dynamic expression of *Oxtr* (29, 30), we looked at the expression of *Magel2* and *Oxtr* transcripts in the anterior hippocampus at P7 and P28, using RNAscope technique. At P7, we detected *Oxtr* and *Magel2* mRNAs in the aCA2/CA3d region with *Magel2* more expressed in the deep layer of the *stratum pyramidale* (Figure 2A). At P28, the level of *Magel2* transcripts was reduced but still present in the deep layer of aCA2/CA3d region and *Oxtr* transcripts were also strongly expressed in pyramidal cells. Expression of *Magel2* and *Oxtr* was also detected in few cells of the *stratum oriens* and *stratum radiatum* where co-expression can be observed. In the DG, an expression of *Oxtr* and *Magel2* was detected in the hilus, with co-localization of both transcripts. Then, in parallel, we extracted data from public RNAseq data libraries obtained in adult mice (Allen brain; Linnarson lab: http://celltypes.brain-map.org/rnaseq/mouse/cortex-and-hippocampus; http://mousebrain.org/genesearch.html). It appears that *Oxtr* and *Magel2* are co-expressed in CA3 excitatory neurons (expressing CCK) and also in several interneuron sub-populations expressing SST. Indeed, we observed a co-expression of *Oxtr* and *Sst* mRNAs in hippocampus (Figure 2B).

**Figure 2:**
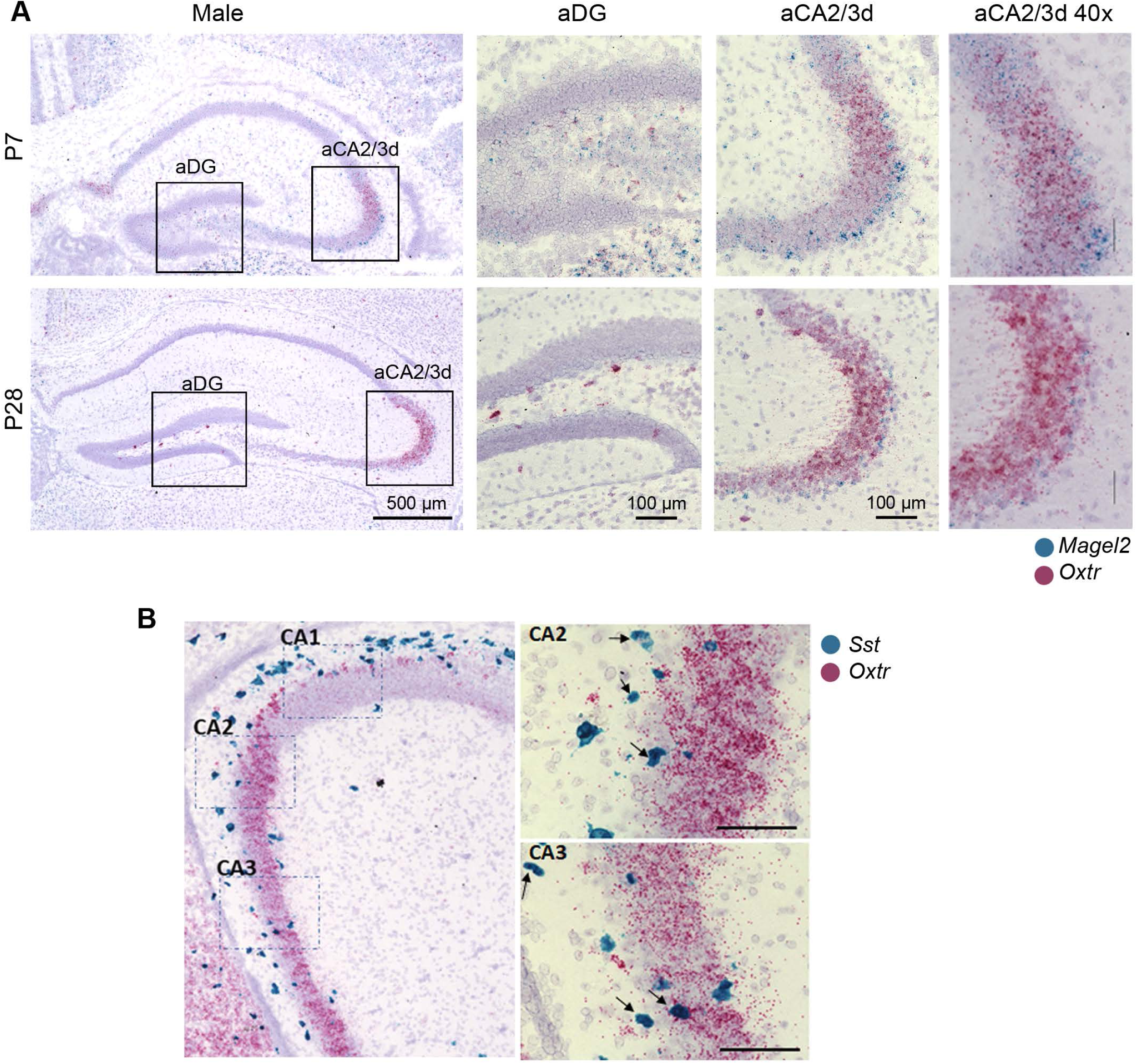
Expression of *Magel2* and *Oxytocin receptor* (Oxtr) transcripts in hippocampus of wild-type male mice at P7 and P28. (**A**) Representative image obtained by RNAscope technology showing the respective localization of *Magel2* (blue) and *Oxtr* (pink) transcripts in dentate gyrus (DG) and aCA2/CA3d region of hippocampus. (**B**) Representative image obtained by RNAscope technique showing the respective localization of *Sst* (blue) and *Oxtr* (pink) transcripts in the aCA2/CA3 region from hippocampal slices of WT male pups at P10. Arrows indicate colocalization of both transcripts in the same cell. Scale bar: 100µm.

We therefore tested the hypothesis that the aCA2/CA3d and DG regions are involved in the social memory deficit of *Magel2-KO* mice.

### Social memory test activates aDG and aCA2/CA3d in WT and *Magel2-KO* mice

WT and *Magel2-*KO mice were sacrificed 90 min after the end of social memory test (+SI, for Social Interactions) or without being tested (-SI) and their brains examined for cFos immunolabeling, a marker of neuronal activity, in the aDG and aCA2/CA3d regions (Figure 3A,B). WT-SI and *Magel2-*KO-SI mice showed a similar quantity of cFos positive cells in both regions. In the aCA2/CA3d region, WT+SI versus WT-SI (Figure 3C,D) showed a significant increase (x1.8) in the number of cFos+ cells; an increase (x2.2) was also observed in *Magel2-* KO+SI mice compared with *Magel2-*KO-SI (Figure 3C,E), notably a 23% significant increase of cFos activated cells was observed in *Magel2-*KO+SI compared with WT+SI (Figure 3D). In the aDG, mainly in the hilus and *stratum granulare,* a significant increase of ∼60% of cFos+ cells was observed in both WT+SI and *Magel2-*KO+SI compared with untested (-SI) mice. Overall, these data confirm a strong activation of neurons in aDG and aCA2/CA3d regions following social memory test in both WT and *Magel2-*KO mice, with an increased activation in the aCA2/CA3d *Magel2-*KO region.

**Figure 3:**
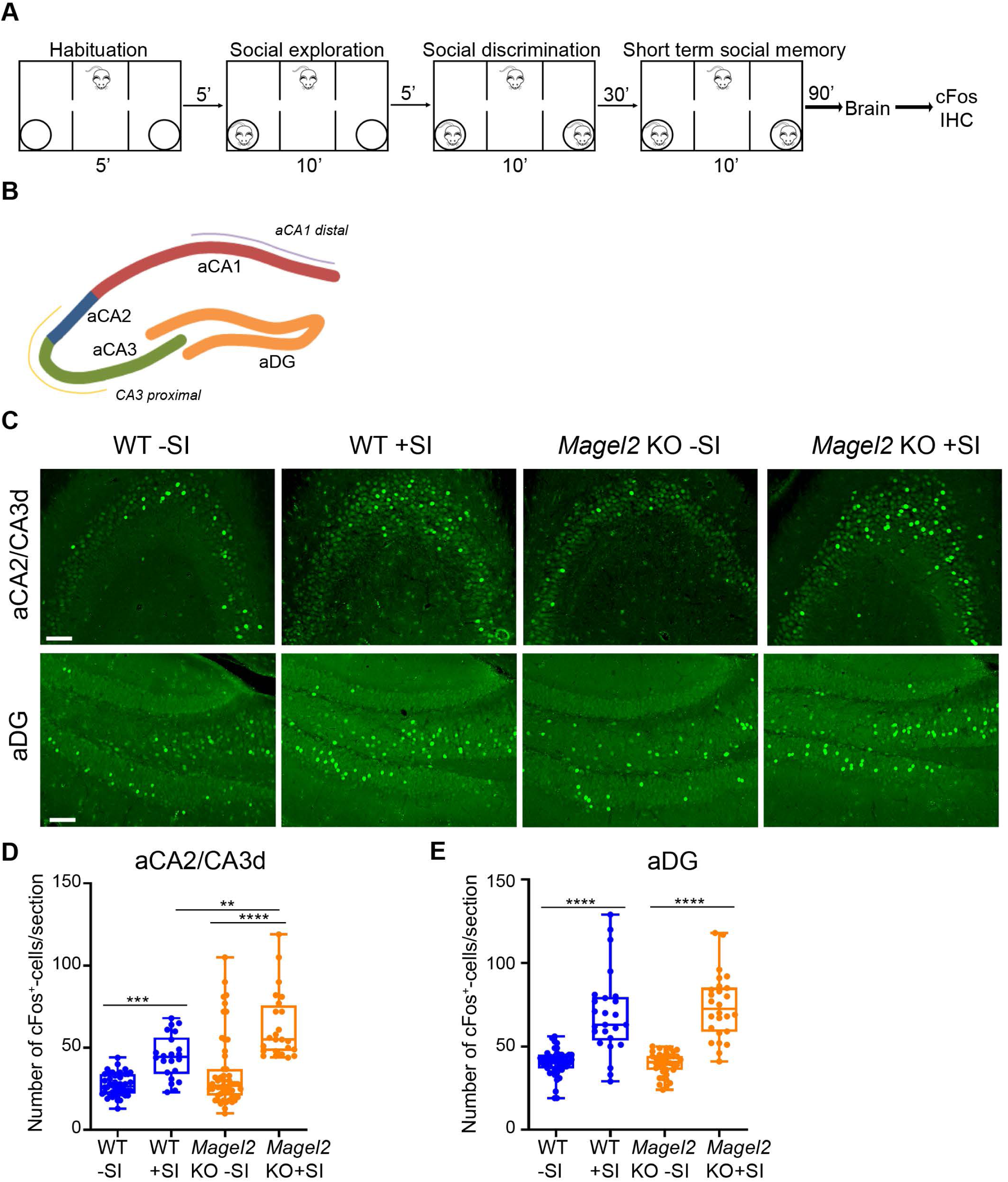
cFos activity in aCA2/CA3d and aDG regions of *Magel2* KO and WT male mice following the social memory task in the three-chamber test. (**A**) Paradigm of the three-chamber test (+SI) followed 90 min later by dissection of the brains and immunohistochemistery experiments. Control mice (-SI) were not tested in the three-chamber test. (**B-C**) cFos-immunolabeling on coronal brain sections in the aCA2/CA3d and aDG regions as indicated in (B) of WT-SI, *Magel2* KO-SI, WT+SI and *Magel2* KO+SI mice (**C**). (**D-E**) Quantification of cFos+ cells/section in WT-SI (n=46, N=6), WT+SI (n=24, N=4), *Magel2* KO-SI (n=38, N=4) and *Magel2* KO+SI (n=24, N=4) in the aCA2/CA3d region (**D**) and in the aDG region (**E**). N: number of animals, n: number of sections/hippocampus. Scale bar: 500µm (B); 100µm (C). Data represented in box and whisker-plots report the number of cFos + cells by sections (6 sections/ hippocampus) with the median (Q2) and quartiles (Q1, Q3) for the genotype and treatment. One-way ANOVA + Dunnett’s *post hoc* test, ***P<0.001. Statistical analysis is reported in Supplemental Table 2.

### High levels of OT-binding sites in *Magel2-*KO hippocampi are reduced by neonatal OT-treatment

We then looked at the distribution of OT-binding sites, reflecting the presence of functional OXTRs, in *Magel2-*KO-vehicle or *Magel2-KO*+OT compared with WT-vehicle hippocampi by autoradiography (Figure 4). We observed a significant increase of OT-binding sites in the aCA2/CA3d (100%, Figure 4A,B) and aDG (80%, Figure 4C,D) regions but not in the ventral region (Figure 4E,F). In *Magel2-KO*+OT we observed a normalization of the amount of OT-binding sites in the aDG, this amount is also decreased in the aCA2/CA3d region but remains high compared to the WT (Figure 4A,B). This binding study indicates subregions specific modulation of OXTRs in the hippocampus of *Magel2*-KO compared with WT. We then wondered if this specific effect could be linked to changes in neuronal subpopulations in these sub-regions.

**Figure 4:**
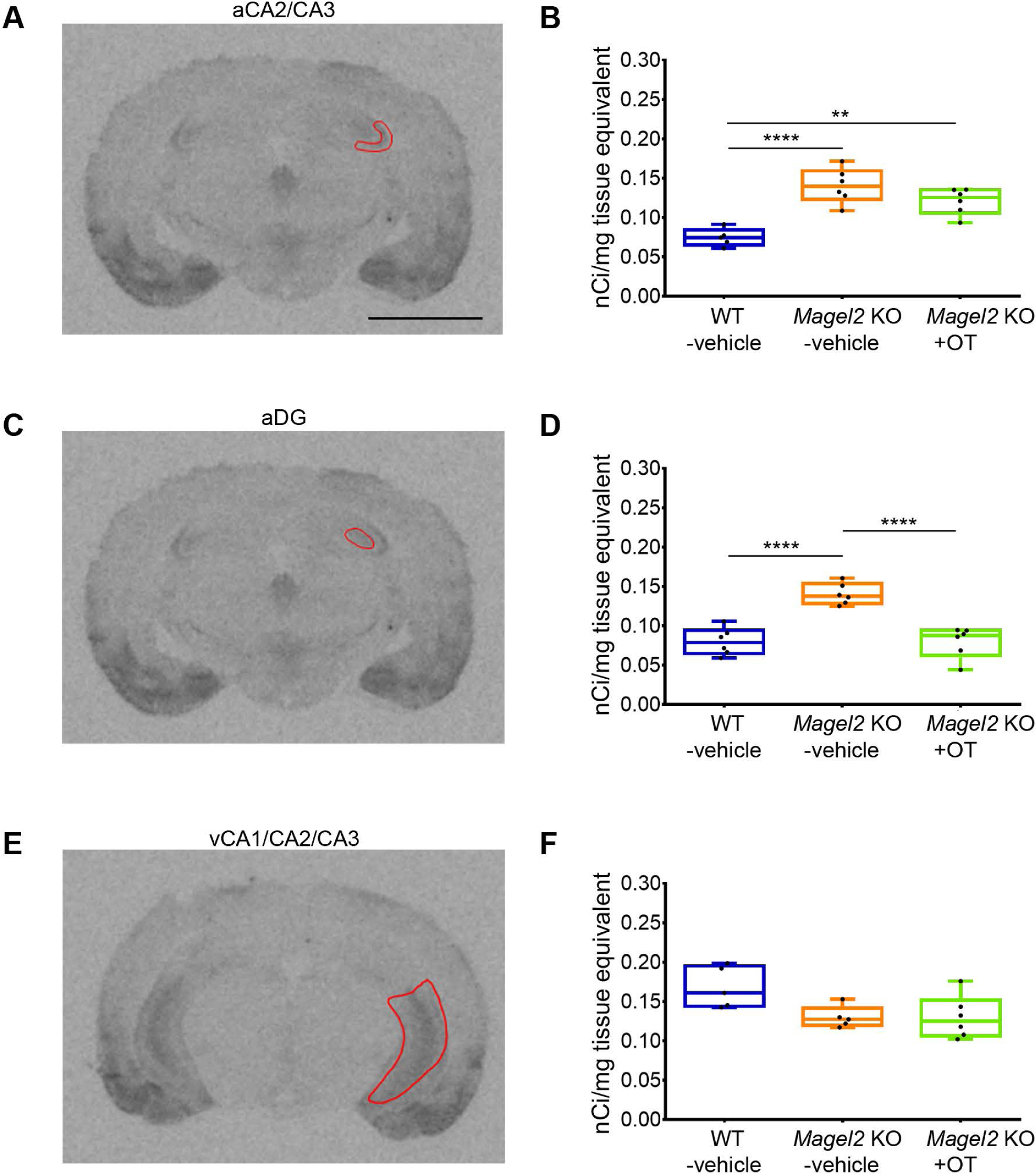
Quantification of OT binding sites by brain autoradiography in *Magel2* KO male mice treated with OT or vehicle versus WT-vehicle male mice. (**A-C-E**) Representative sections of autoradiographic labeling of OT binding sites displayed in grayscale, showing the regions of interest (ROI) selected for analysis: (**A**) anterior CA2/CA3 (aCA2/CA3), (**C**) dentate gyrus (aDG) and (**E**) ventral CA1/CA2/CA3 (vCA1/CA2/CA3) regions of hippocampus. (**B-D-F**) Quantification of OT binding sites expressed as nCi/mg of tissue equivalent in (**B**) anterior CA2/CA3, (**D**) dentate gyrus and (**F**) ventral CA1/CA2/CA3 regions of hippocampus. Histograms report median (Q2) and quartiles (Q1, Q3). OT binding sites in nCi/mg of tissue equivalent. 3 mice and 6 hippocampi have been analyzed for each group. Data represented in box and whisker-plots report the quantity of radiolabeling by hippocampus, with scattered plots that show individual data points. One-way ANOVA + Bonferroni *post hoc* test, **P<0.01, ****P<0.0001. Scale Bar: 3 mm. Statistical analysis is reported in Supplemental Table 4.

### An increased number of SST+ neurons in the aCA2/CA3d and aDG regions of *Magel2-*KO adult mice is normalized by neonatal OT-treatment

In the anterior adult hippocampus OXTRs are expressed in pyramidal cells of aCA2/CA3d region and mainly in SST and/or PV interneurons of aCA2/CA3d and aDG (23, 27). In *Magel2-*KO adult mice, the number of SST+ cells was higher than in WT mice in both aCA2/CA3d (more 60%) and aDG (more 80%) regions (Figure 5 A-F and M-N). After a neonatal OT-treatment in *Magel2-*KO, the number of SST+ cells was slightly but significantly decreased in both aCA2/CA3d (less 12%) and DG regions (less 17%) mice compared with untreated WT mice (Figure 5 G-L and O-P). OT-treatment of *Magel2-*KO pups has resulted in a significant decrease of SST+ neurons in adult mutant compared with WT hippocampi. PV+ cells were equally abundant in both genotypes (Figure 5-Supplement1). We then expected that this change in the number of SST+ cells had consequences in the alteration of the excitation/inhibition (E/I) balance, an electrophysiological feature frequently associated with multiple neurodevelopmental disorders.

**Figure 5:**
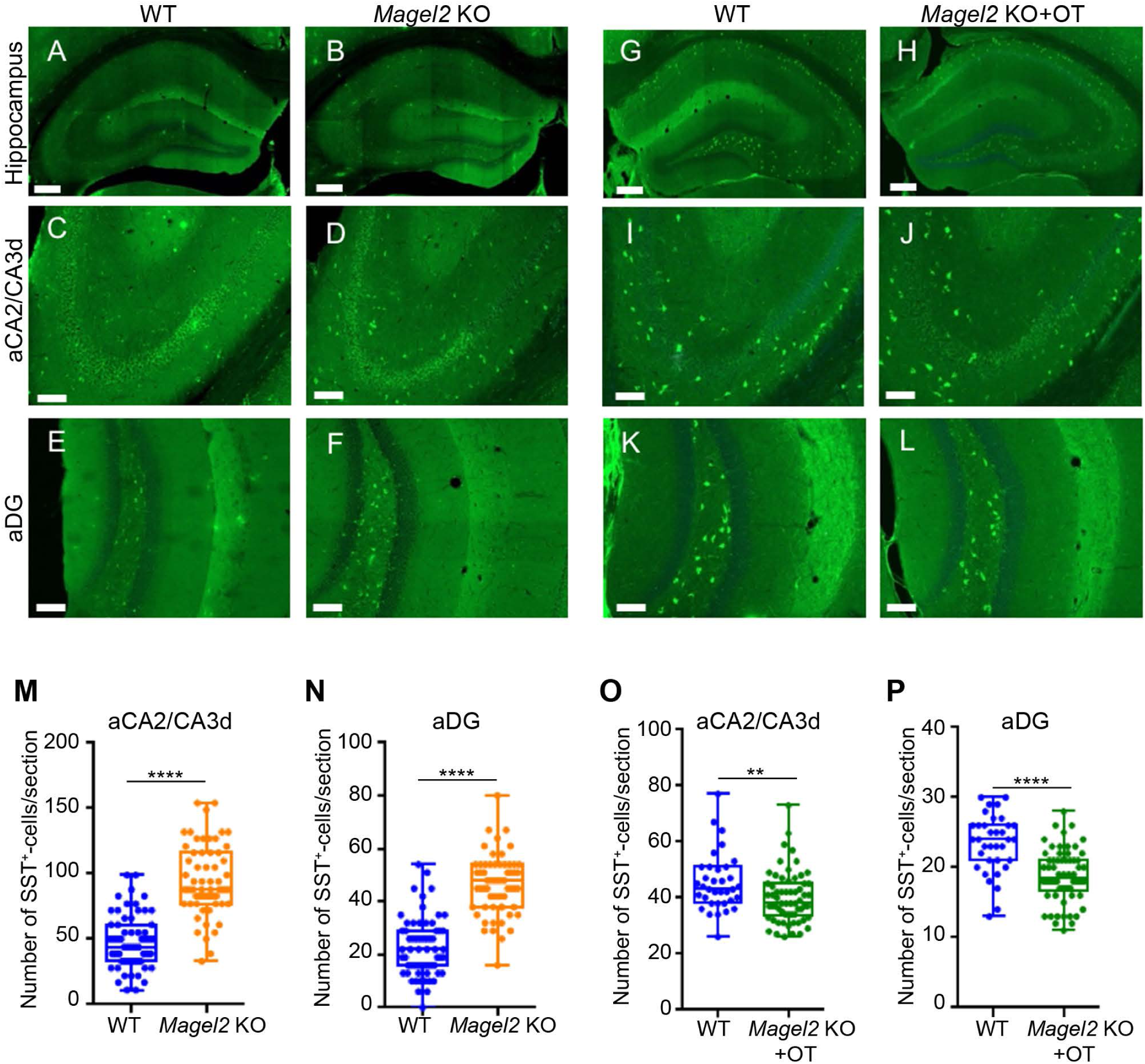
Quantification of somatostatin (SST) immunopositive cells in the anterior hippocampus region of *Magel2* KO adult mice having or not been treated by OT in the first week of life and compared with WT mice. (**A-L**) Immunolabeling on coronal hippocampal sections at adulthood in WT (A,C,E) and *Magel2 KO* (B,D,F), and in WT (G,I,K) and *Magel2 KO*+OT (H,J,L) with a magnification in the aCA2/CA3d region ( C,D,I,J) and in the DG region (E,F,K,L) in which the SST+ cells are counted. (**M-P**) Number of SST+ cells by section in both aCA2/CA3d (M,O) and aDG (N,P) and comparing WT (N=4, n= 48) with *Magel2* KO (N=4, n=48) animals (M,N) or WT (N=3, n=36) with *Magel2* OT+ (N=5, n=56) (O,P) mice. N: number of animals, n: number of sections/hippocampus. Data represented in whisker-plots report the number of SST+ cells by section with Q2(Q1, Q3), with scattered plots showing individual data points. Mann-Whitney Test **P<0.01, ***P<0.001. Scale bar (A-H): 500 µm; (C-L): 100 µm. Statistical analysis is reported in Supplemental Table 5.

### Neonatal OT-treatment normalizes the increased GABAergic activity in *Magel2-KO* aCA2/CA3d neurons but reduces the glutamatergic activity in both mutant and WT mice

Hippocampal brain slices of WT, *Magel2-KO*, WT+OT and *Magel2-KO*+OT male mice were analyzed using whole cell patch clamp to record the activity of aCA2/CA3d pyramidal neurons (Figure 6A). Spontaneous activities analysis (Figure 6B) revealed a reduced amplitude of postsynaptic glutamatergic currents (sGlut-PSCs) in *Magel2-KO* as compared to WT mice (x1.7 less, Figure 6D), while the frequency of sGlut-PSCs was not changed (Figure 6C). The same *Magel2-KO* neurons presented a significant increase in GABAergic (sGABA-PSCs) frequency (x1.8, Figure 6E) while the amplitude of sGABA-PSCs was similar to that of WT (Figure 6F). Patch clamp recordings of the glutamatergic and GABAergic miniature currents (mGlut-PSCs and mGABA-PSCs, respectively) showed a significant reduction in amplitude of mGlut-PSCs, with no change in their frequency, in *Magel2-*KO neurons compared to WT (x1.4 less, Figure 6-Supplement1B-C), but no differences in frequencies and amplitudes of mGABA-PSCs (Figure 6-Supplement1D-E). An abnormal dendritic morphology of the pyramidal neurons could be associated with the alterations of the neuronal activities. However, we showed that dendritic morphology of *Magel2-*KO and WT CA3 pyramidal neurons were similar (Figure 6-Supplement2). Altogether, those results show that, in *Magel2-*KO pyramidal neurons of aCA3d, there is a significant increase in the GABA/Glutamate ratio with no change in their neuronal morphology.

**Figure 6:**
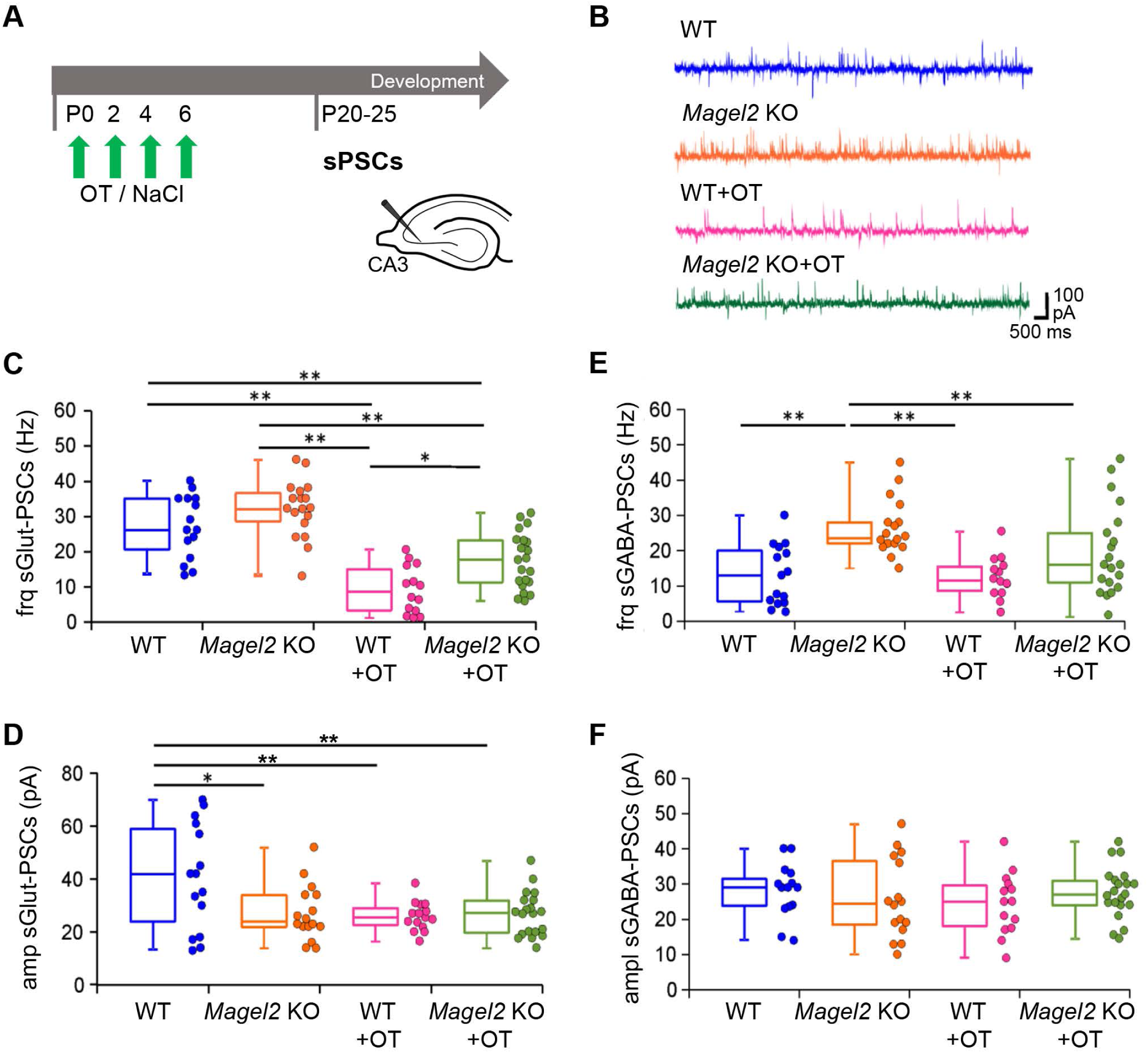
Spontaneous Glutamatergic and GABAergic synaptic activity of CA3 pyramidal neurons in the anterior hippocampus region of *Magel2* KO mice versus WT juvenile mice and having been OT-treated or vehicle-treated in neonates. (**A**) Paradigm of the test. WT or *Magel2 KO* mice have been or not injected with OT in the first week of life then neurons are recorded in brain slices at P25. (**B**) Examples of whole cell recordings performed at a holding potential of −45 mV for each genotype or treatment. The glutamatergic synaptic currents are inwards and the GABAergic synaptic currents are outwards (**C-D**) Values in the different genotypes and treatment of the Glut-sPSCs frequency (C) and amplitude (D). (**E-F**) Values in the different genotypes and treatment of the GABA-sPSCs frequency (E) and amplitude (F). *Magel2* KO (N= 7, n=16), WT (N=7, n=15), *Magel2* KO+OT, (N=5, n=21) and WT+OT (N=4, n=15) have been analyzed, with N: number of mice and n: number of recorded cells. Data represented in whisker-plots report the different values of recorded cells with mean ±SEM, with scattered plots showing individual data points. One-way ANOVA + Tuckey *post hoc* test. *P<0.05, **P<0.01. Statistical analysis is reported in Supplemental Table 6.

We next investigated the effects of OT-treatment on the GABA/Glutamate balance in WT and *Magel2-*KO neurons. Quite unexpectedly, the frequency (x3 less) and amplitude (x1.7 less) of sGlut-PSCs were significantly reduced in WT+OT mice compared with WT-vehicle (Figure 6C,D). In *Magel2-*KO mutants, OT-treatment did not modify the amplitude but reduced the frequency (x2.9 less) of sGlut-PSCs compared with *Magel2-*KO mice (Figure 6C,D). The GABAergic activity (amplitude and frequency) was not changed in WT mice after an OT-treatment (Figure 6E,F). In *Magel2-*KO, OT-treatment decreased (x1,9 less) significantly the frequency of sGABA-PSCs restoring a frequency similar to WT (Figure 6E); no effect was observed on the amplitude of sGABA-PSCs (Figure 6F). These results show that OT administration in the first week of life normalized the frequency of spontaneous GABAergic activity in *Magel2-*KO neurons. Neonatal OT-treatment reduced strongly the frequency of glutamatergic activity in *Magel2-*KO and the reduction is even stronger on the amplitude and frequency in WT mice. Notably, WT mice had normal behaviors (Figure1-Supplement3).

### A delay of the excitatory-to-inhibitory developmental GABA-shift in *Magel2-KO* hippocampal neurons is corrected by neonatal OT-administration

Because *Oxtr* and *Magel2* are co-expressed in aCA2/CA3d hippocampus in infancy (P7), and because in *Oxtr* KO mice (35), as in several models (38) of autism, the depolarizing to hyperpolarizing developmental GABA shift is delayed, we investigated the GABA shift timing in *Magel2-KO* pups. First, we performed calcium (Ca^2+^) imaging experiments by measuring the percentage of neurons showing GABA-induced Ca^2+^ responses in developing hippocampal neuronal cultures of *Magel2-KO* and WT embryos collected at E18.5 d.p.c (DIV0 for days in vitro 0) (Figure 7A,B). At DIV4, we found a significant two-fold higher proportion of *Magel2-KO* neurons increasing Ca^2+^ upon GABA stimulation compared with WT. This difference was abolished at DIV8 and DIV11 with a marked decrease of the percentage of responsive neurons in both genotypes. In *Magel2-*KO, KCl-stimulation did not increase Ca^2+^ responses at DIV2, DIV4 and DIV8 compared to WT (Figure 7-Supplement1A), suggesting that the voltage-operated calcium channels were not responsible for the higher proportion of *Magel2-*KO GABA evoked neuronal responses upon stimulation. Altogether, these results showed a developmental delay in GABA-induced Ca^2+^ responses in cultures of *Magel2-*KO embryonic hippocampal neurons and suggest a developmental GABA-shift delay in *Magel2-*KO hippocampal neurons.

**Figure 7.**
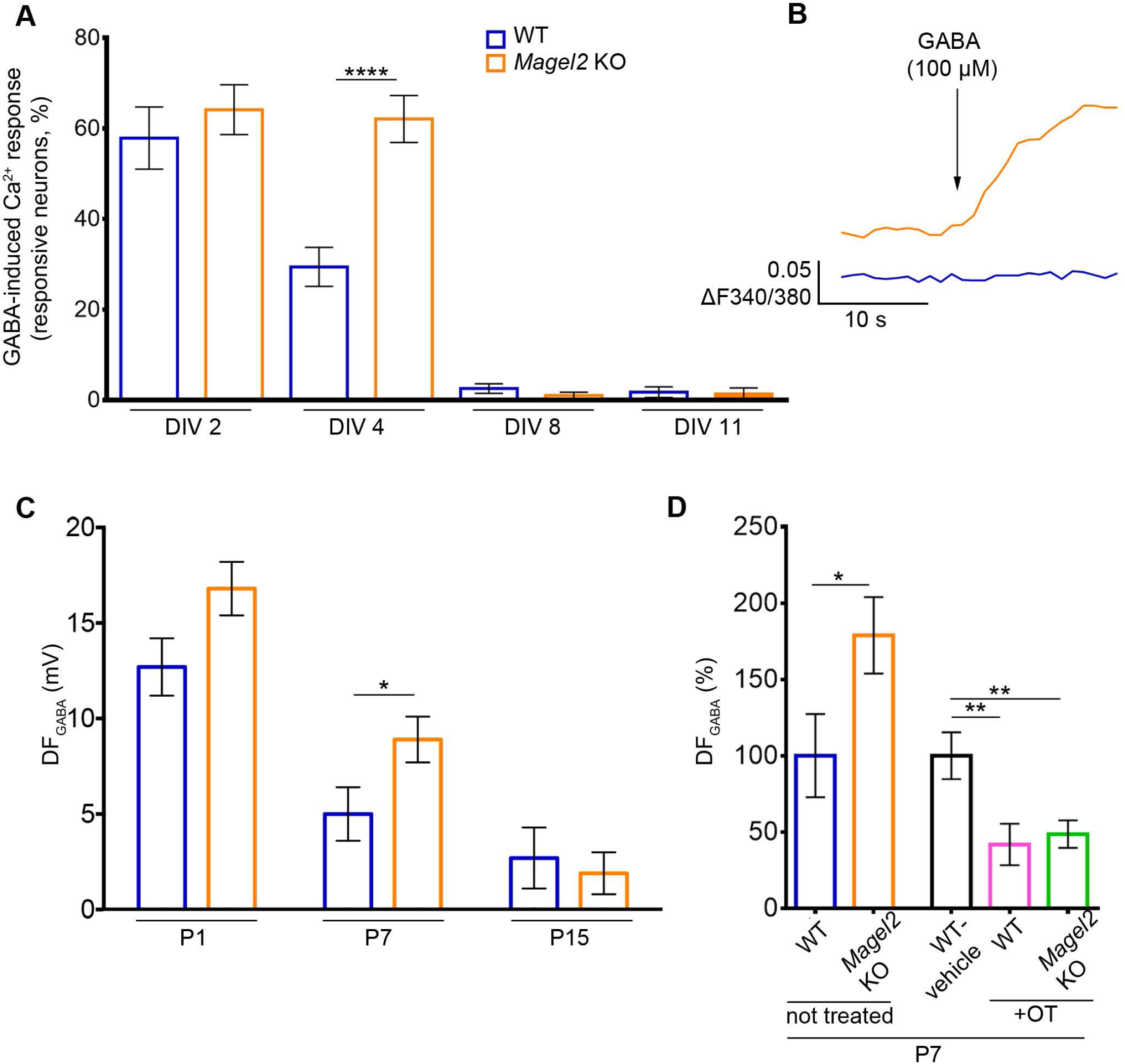
The excitatory-to-inhibitory developmental GABA-shift in *Magel2* KO versus WT hippocampi and the effect on an OT-treatment. (**A-B**) GABA-induced Ca^2+^ responses in *Magel2* KO developing hippocampal neuronal cultures versus WT. (**A**) Percentage of WT and *Magel2* KO E18 hippocampal neurons showing GABA-induced Ca^2+^ responses at selected in vitro days (DIV). (**B**) Representative traces of [Ca^2+^]i variations (delta F340/380) in DIV4 WT and *Magel2* KO neurons upon 100 μM GABA administration. Data are presented in histograms with mean ± SEM; unpaired t test with Welch’s correction: ****P<0.0001. (**C-D**) Measures of the driving force for GABA (DF_GABA_), using cell-attached recordings of single GABA_A_ channels, of aCA3d pyramidal neurons. (**C**) Average values of the DF_GABA_ at P1, P7 and P15 in *Magel2* KO (P1 N= 3, n= 20; P7: N= 7, n= 56; P15: N= 4, n= 29) versus WT (P1 N= 3, n= 19; P7: N= 6, n= 42; P15: N= 3, n= 23) mice. Data are presented in histograms with mean ± SEM; unpaired t test with Welch’s correction: *P<0.05. (**D**) Graph reporting the relative changes of the DF_GABA_ at P7 in untreated *Magel2*-KO mice (N= 7, n= 56) compared with WT mice (N= 6, n= 42) and in OT-treated WT (N= 3, n= 37) and *Magel2*-KO (N= 4, n= 56) mice compared with WT-vehicle (N= 3, n= 37) mice. N: number of mice and n: number of recorded cells. One-way ANOVA + Dunnett’s *post hoc* test: **P<0.01. Statistical analysis is reported in Supplemental Table 7.

Cell-attached recordings of single GABA_A_ channels/receptors were then performed in acute brain slices in order to measure the Driving Force of GABA_A_ (DF_GABA_) and therefore the hyperpolarizing or depolarizing response of GABA under conditions where intracellular chloride levels are not altered by the recording pipette. At P1, we observed a tendency to an increase of DF_GABA_ in *Magel2-*KO compared with WT. At P7, the DF_GABA_ was significantly increased (x1.8) in mutant neurons. This difference was abolished at P15. Since *Magel2* and *Oxtr* are expressed in interneurons (INs), in which a GABA-shift has been also described (39), we also measured the DF_GABA_ in INs and observed similar values in CA3 interneurons of mutant and WT mice (Figure 7-Supplement1C). At P7, the resting membrane potential (Figure 7-Supplement1D), the conductance (Figure 7-Supplement1F) and capacitance (Figure 7-Supplement1E) did not differ statistically between WT and *Magel2*-KO neurons. Altogether these data suggest a transient higher GABA depolarizing activity at P7 in CA3 pyramidal neurons of *Magel2-KO* pups and consequently a delay in GABA-shift. Then, at P7, we assessed the effect of the neonatal OT-administration in *Magel2-KO* and WT neonates compared with WT-vehicle pups. Both *Magel2-KO*+OT and WT+OT mice showed a significant decrease (x2.4 less and x2 less, respectively) in the DF_GABA_ values compared with WT-vehicle (Figure 7D), suggesting a reduction in GABA depolarizing activity following a neonatal OT-administration.

### Post-translational changes in cation-chloride co-transporter KCC2 in *Magel2-KO* hippocampus

We then asked if the altered DF_GABA_ values could be due to alteration in the expression of the neuronal transporters of Cl^-^in *Magel2-KO* mice. The neuronal level of [Cl^-^]_i_ and Cl-dependent depolarizing or hyperpolarizing strength of GABA are determined by complex mechanism involving primarily Cl^-^ extrusion by potassium/chloride cotransporter type 2 (KCC2) whose expression increases progressively during neuronal maturation (40). In developing WT hippocampal neurons, the emerging activity of KCC2 contributes to progressive lowering of [Cl^-^]_i_ that at P7 shifts GABA action from depolarizing to hyperpolarizing. As consequence, the activation of GABA_A_R produces neuronal Cl^-^ influx.

Quantitative western blot analysis of the total KCC2 protein expression in hippocampi of P7 mice did not reveal statistically significant difference of the amount of KCC2 between WT and *Magel2-KO* animals (Figure 8A,B). However, the ion-transport activity of KCC2 and its stability at the cellular plasma membrane depend on post-translational modifications of multiple phosphorylation sites (41). We therefore used phospho-site-specific antibodies, previously shown to quantitatively monitor changes in KCC2 phosphorylation (42–44). Currently, a limited number of such phospho-specific antibody is available. They are directed against the well-known KCC2’s phospho-sites Ser^940^ (45) and Thr^1007^ (43, 44). Western blot analysis revealed that the *Magel2-KO* hippocampi (as compared to WT) were characterized by significantly decreased amount of the phosphorylated form of Ser^940^ (P-Ser^940^). The amount of phosphorylated Thr^1007^ (P-Thr^1007^) was not statistically different, although a small but not significant increase was observed in *Magel2-KO* mice (Figure 8A,B). At P7, the decreased P-Ser^940^/P-Thr^1007^ ratio in *Magel2-KO* mice may thus result in predominance of KCC2 internalization over surface expression. As a consequence of the decreased amount of surface expressed molecules, the Cl^-^ extrusion ability of KCC2 is decreased, causing an increase of [Cl^-^]_i_ and could induce a depolarizing shift of GABA described above (Figure 8C).

**Figure 8.**
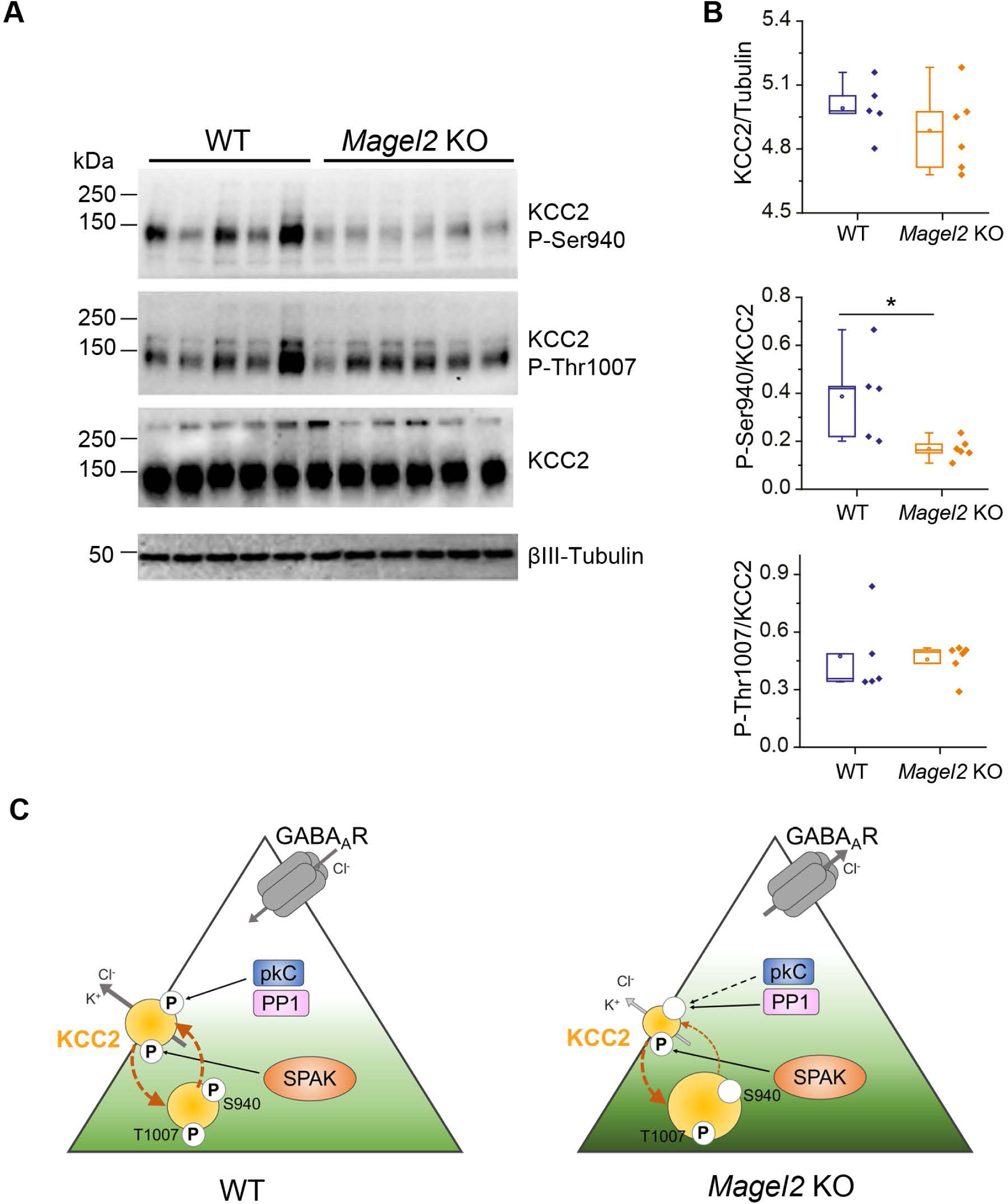
Abundance and phosphorylation state of KCC2 in WT and *Magel2* KO pups. (**A**) Immunoblot analysis of WT (N=5) and *Magel2* KO (N=6) hippocampi of P7 mice with pan-KCC2 antibody or phosphorylation site-specific antibodies recognizing P-Ser^940^ or P-Thr^1007^ of KCC2. An antibody recognizing neuron-specific ß3 tubulin was used to normalize the quantity of proteins. Numbers on the left indicate molecular weight. (**B**) Boxplots report band intensities from (A) as Q2(Q1,Q3), with scattered plot showing individual data points. Mann-Whitney test, *P<0.05. (**C**) A model of KCC2-dependent control of neuronal Cl^−^ in *Magel2* KO pups. At this stage of neuronal development, the surface expression of KCC2, that determines its ion-transport activity, depends on the ratio of reciprocal phosphorylation of its Ser^940^ and Thr^1007^ residues. The Ser^940^ phosphorylation increases KCC2’s cell surface stability, whereas the Thr^1007^ phosphorylation exerts opposite to Ser^940^ effect and favors internalization (shown with brown arrows). Compared to WT, the CA3 neurons in hippocampi from *Magel2* KO mice are characterized by depolarizing action of GABA (e.g. activation of GABA generates Cl^-^ efflux) that reflects higher [Cl^-^]_i_. In *Magel2* KO hippocampi the amount of KCC2’s Ser^940^ phosphorylation is significantly lower as compared to WT hippocampi whereas the amount of phosphorylated Thr^1007^ remains unchanged. Respectively, the decreased P-Ser^940^/P-Thr^1007^ ratio results in predominance of KCC2 internalization over surface expression. As consequence of the decreased amount of surface expressed molecules, the Cl^-^ extrusion ability of KCC2 is decreased that causes increase of [Cl^-^]_i_ and depolarizing shift of GABA. The model includes also important components that are known to control the level of Ser^940^ and Thr^1007^ phosphorylation. The Ser^940^ is directly phosphorylated by kinase C (pkC) and dephosphorylated under pathology conditions by protein phosphatase type 1 (PP1). The Thr^1007^ is directly phosphorylated by SPAK. It remains to be elucidated whether in *Magel2* KO mice the decreased level of Ser^940^ results from reduction of pkC activity or enhancement of PP1 action. Statistical analysis is reported in Supplemental Table 8.

## DISCUSSION

Here we investigated, in hippocampus, the mechanisms by which peripheral administration of OT in neonates acts to restore normal social memory in *Magel2-KO* mice. Peripheral OT-administration in neonates permanently rescued almost all the hippocampal alterations (quantity of OT-binding sites, number of SST-positive interneurons and an increase in the GABAergic activity of pyramidal neurons) that we have characterized and that are associated with the increase of GABAergic activity observed in *Magel2-KO* adult mice; but a decrease of glutamatergic activity is still present. Those alterations are related with the OT-signaling pathway and relevant to explain the loss of social memory. However, a significant impact of OT-treatment, reducing the glutamatergic activity, was also observed in wild-type mice but all performed behavioral tests were normal. A significant effect of OT-administration on the delayed excitatory-to-inhibitory developmental GABA-shift (at P7), delays that are observed in various neurodevelopmental disorders, underlies the therapeutical use of neonatal oxytocin in these diseases.

### The effect of peripheral OT-administration in *Magel2-KO* neonates

Here, with social memory study, in addition to our previous work, we have shown long-term and beneficial effects of a peripheral administration of OT in *Magel2-KO* neonates that rescues nearly all social and cognition deficits described in adult *Magel2-KO* mice (14). However, previously, we did not investigate neither the neurobiological causes of these deficits nor the effects of OT on those alterations. We focused this study on social memory because the mechanisms by which OT controls social memory via the OXTR-expressing neurons in hippocampus are known. We clearly demonstrated that 1) the neurobiological alterations found in *Magel2-KO* mice involve the hippocampal brain OT-circuitry and 2) peripheral administration of OT in neonates impacts this circuitry. Thus, those results give a clear and positive response to the debated question on the action of peripheral administration of OT on the brain. Is it a direct or indirect effect? It might be both. Today, several studies converge to propose that peripheral OT goes through the Brain Blood Barrier via an active mechanism (RAGE transporters) and/or a passive mechanism, in neonates, when this barrier is more permeable. The observed long-lasting OT effects could result from a strong impact of OT administration in key developmental hippocampal processes such as the developmental GABA-shift (as discussed below) and can also be achieved by epigenetic modifications that impact gene expression such as the *Oxtr* expression, as observed in prairie voles following a maternal OT administration (46). Transcriptomic and proteomic studies at different developmental ages would help to understand the life-long effect of an early OT-treatment in mutant and WT mice.

### The lack of *Magel2* alters the OT-system: causes and consequences

*Magel2* is expressed in hypothalamic OT neurons and, in *Magel2*-KO neonates, we observed a deficit in the quantity of the mature form of OT although we detected an accumulation of the intermediate non-mature forms of OT suggesting a problem of hormonal processing (16). Here we showed a co-expression of *Magel2* and *Oxtr* transcripts in the hippocampal neurons. Thus, given the role of *MAGEL2* in ubiquitination, actin regulation and endosomal sorting processes (47), the absence of *Magel2* expression could induce post-translational modifications of various processes in OT and OXTR expressing neurons, suggesting that dysregulation of the OT-system in *Magel2-KO* mice goes beyond OT expression.

Here we showed that the excitatory-to-inhibitory GABAergic developmental sequence is transiently delayed in *Magel2*-KO pyramidal neurons during the first week of life, with GABA_A_-mediated responses more depolarizing in *Magel2-KO* that WT mice at P7. We further showed that this electrophysiological deficit corresponds well with decreased functional KCC2 at the cell membrane, caused by a deficit of KCC2 phosphorylation (on Ser^940^). Notably, Ser^940-^ phosphorylation is controlled by OXTR activation via a PKC-dependent pathway and allows translocation of KCC2 to the cell membrane (35), enhancing KCC2 mediated Cl^-^ transport (45, 48). This mechanism is relevant to control the GABAergic developmental sequence *in vitro* and possibly *in vivo* (49).

The functional consequences of this delayed developmental GABA-shift in *Magel2-KO* pyramidal neurons are not clearly established. However, there are compelling reasons to suspect that transient disruption of GABAergic maturation in the immediate postnatal period might be sufficient to permanently alter neural circuit dynamics. Indeed, P7 is a critical milestone in the development of GABAergic neurons in the mouse neocortex and hippocampus, characterized by major changes in network dynamics (e.g. end of *in vitro* giant-depolarizing-potentials (50) and *in vivo* early sharp waves (51)), intrinsic membrane properties (e.g. input resistance) and synaptic connectivity (52). Altogether those data suggest that the absence of *Magel2* delays neuronal maturation during this critically vulnerable period of brain development, resulting in a distinct adult phenotype. Whether the delay of the GABA-shift alone is sufficient to derail neurotypical developmental trajectory remains a key question for future study: notably, similar or longer GABA-shift delays have been observed in several models of autism (53–55) and in *Oxtr* KO mouse models (35). Recently, Kang et al. (56) showed in *Disc1* KO mouse model, that elevated depolarizing GABA signaling is a precursor for the later E/I imbalance (in favor of inhibition) and social impairment. Similarly, we showed that, in a KCC2 mutant mouse, the GABA-shift delay is responsible for the E/I alteration (49).

Importantly, OT-treatment has an opposite action on the excitatory-to-inhibitory GABA-shift with a relative hyperpolarizing effect at P7 in *Magel2-KO* and WT pups compared with WT-vehicle animals. This effect of OT-treatment might modify the maturation of the hippocampal circuitry.

### The E/I ratio and social behavior

Reductions in synaptic signal-to-noise ratio in cortical and hippocampal pyramidal neurons, driven by a change in the ratio of dendritic excitatory and inhibitory synapses, are widely thought to contribute to reduced efficiency of signal processing in ASD, a mechanism known as the E/I ratio hypothesis (57). We confirm E/I imbalance characterized by increased GABAergic activity and lower glutamatergic activity in CA3 neurons in *Magel2-KO* mice, consistent with observations made in some ASD models (58–61). Furthermore, we report that perinatal OT administration restored normal GABAergic activity in *Magel2-KO* mice without improving glutamatergic transmission. Unexpectedly, perinatal OT treatment has a significant impact on the WT neurons inducing a strong reduction of glutamatergic activity without affecting GABAergic activity. This is a significant observation, because it shows that, although the ASD-like behavior *Magel2-KO* animals is correlated with a change in E/I ratio, E/I imbalance in OT-treated WT animals was not sufficient to drive detectable changes in social behavior or cognitive performance. We therefore propose that E/I imbalance characterized by isolated decreased spontaneous glutamatergic transmission is unlikely to underlie the ASD traits investigated here, and suggest that an upper threshold of GABAergic or glutamatergic activity, but not the E/I ratio *per se*, may be important for normal development.

### Role of oxytocin receptors and somatostatin neurons

In adult *Magel2-KO* mice we observed increased OT-binding in the DG and CA2/CA3 regions of the anterior hippocampus compared to WT mice. OT administration in *Magel2-KO* neonates normalized hippocampal OT-binding sites in adulthood, suggesting that the increased expression of OXTR observed in *Magel2-KO* hippocampus may be a consequence of the reduced OT production reported in these animals (16). This observation supports the idea that life-long OXTR expression is to some extent determined by early life OT binding, described as a “hormonal imprinting” effect (46, 62).

Since DG and CA2/CA3 hippocampal OXTRs are expressed in PV and SST interneurons, we quantified those populations and found a significant increase in the number of aDG and aCA2/CA3d SST+-neurons in mutant mice, while the number of PV+-interneurons was not modified. OT-treatment normalized the number of SST-expressing neurons in *Magel2-KO* pups, revealing a causal link between the administration of OT in infancy and the quantity of SST+-neurons. This result may reflect actual changes in the number of SST+-neurons, or alternatively changes in SST expression and hence more reliable detection of SST-synthesizing neurons. Interestingly, OT modulates the activity of the SST+-neurons, increasing the excitability of SST interneurons (63) but no studies report an effect of OT on SST production.

SST interneurons have recently been shown to play a role in the modulation of social behavior (64, 65) and a link between altered social memory and an increase in SST cell number has been recently suggested in LPS-treated female neonates (66). It is tempting to speculate that OXTR-transmission regulates the activity of SST hippocampal interneurons and the production/release of mature SST and impacts social memory. Further work is needed to fully characterize the role of OXTRs on SST interneurons in relation with social memory.

## Conclusions

Oxytocin deficiency, present in the *Magel2-KO* mouse model and in PWS, has also been frequently described in rodent models of ASD (2). Recently, evidences for a unifying role of oxytocin in pathogenic mechanisms responsible for social impairments across abroad range of autism etiologies have been provided (67, 68). Thus, our results demonstrate that peripheral OT-administration in a critical period of time, after birth, represents a viable therapeutic strategy for patients with SYS or PWS and possibly other neurodevelopmental disorders.

## Acknowledgements

The authors thank the Foundation for Prader-Willi Research (FPWR) grants, Agence Nationale pour la Recherche (ANR-14-CE13-0025-01), Thyssen Foundation (Grant 10.16.2.018MN), Prader-Willi France and Fondation Jerôme LeJeune for their financial support.

The authors thank Pr. Simon McCullan, Dr. L Fasano and Dr. C. Gestreau for comments and careful reading of the manuscript. Dr. A. Baude for her technical advices, Dr F. Michel for his help in microscopy and the use of image softwares,

---

**Figure 1-figure supplement 1.**
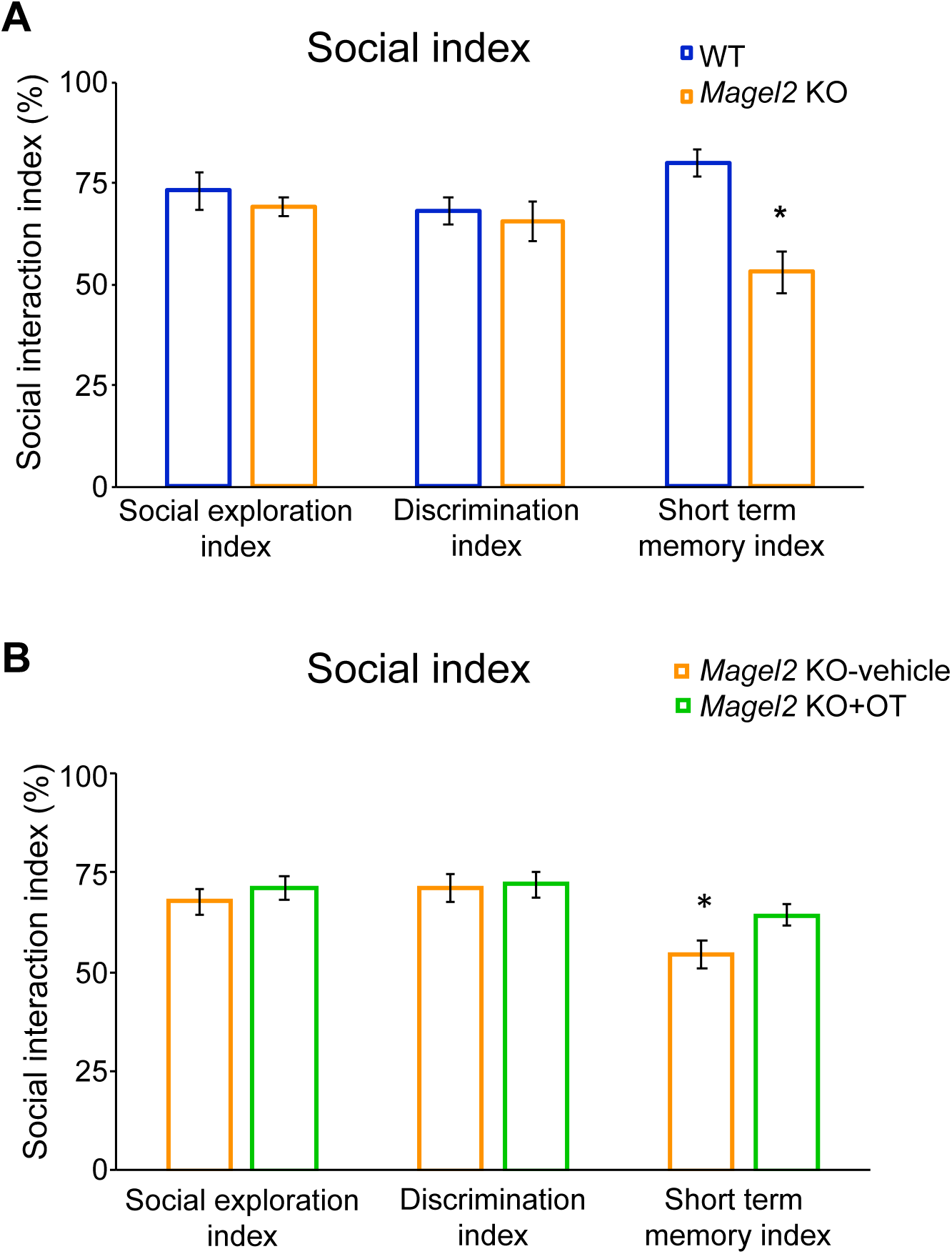
Social-index values comparing Magel2 KO versus WT male mice or Magel2 KO+OT versus Magel2 KO-vehicle male. These indexes report for the social exploration: the sniffing time with S1/ sniffing time with S1 + time in empty room x 100; for the discrimination: the sniffing time with S2/ sniffing time with S1 + sniffing time with S2 time x 100 and for short term memory: the sniffing time with S3/ sniffing time with S1 + sniffing time with S3 time x 100. They are measured in (A) Magel2 KO versus WT mice and (B) Magel2 KO+OT versus Magel2 KO-vehicle. It appears clearly that social exploration and social discrimination are similar between WT and Magel2 KO (A) and between Magel2 KO-vehicle and Magel2 KO+OT (B). Only short memory index is decreased in Magel2 KO compared with WT and increased in Magel2 KO+OT compared with Magel2 KO-vehicle. Data in histograms report social indexes calculated for each individual as mean ± SEM. Mann-Whitney test, *P<0.05. Statistical analysis is reported in Supplemental Table 1-Supplement 1.

**Figure 1-figure supplement 2.**
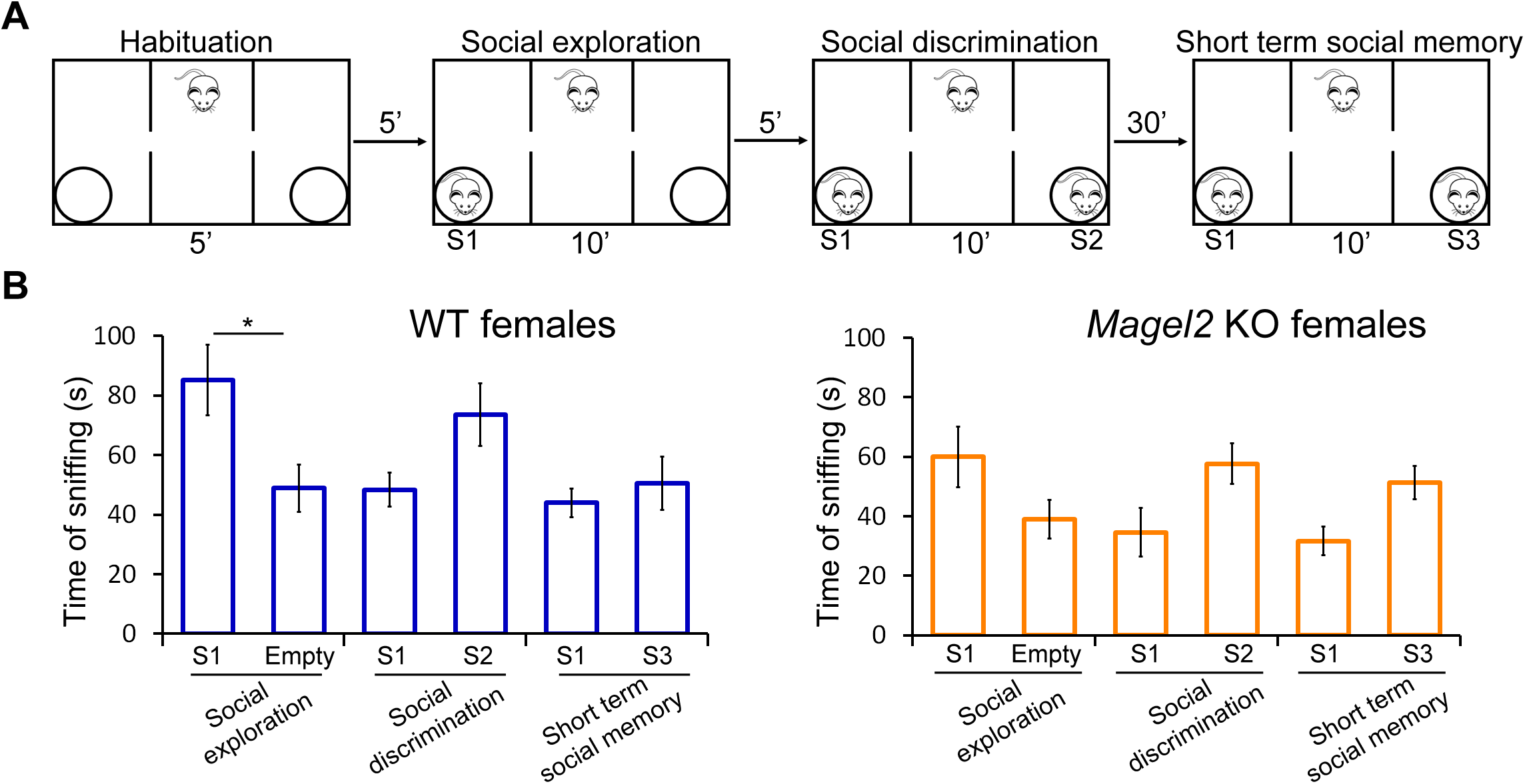
Social behavior in three-chamber test of female Magel2 KO adults versus WT adults. (A) Paradigm of the three-chamber test. Sniffing time between mice is measured in each test. (B) WT (N=11) females do present differences in social exploration but do not present significant differences in social discrimination and short-term social memory, suggesting that the three-chamber test is not relevant to assess the social behavior in females. Similarly, Magel2 KO (N=9) females do not present significant differences in all three steps of the paradigm. Data in histograms report interaction time (time of sniffing in seconds) as mean ± SEM. Mann-Whitney test, *P<0.05. Statistical analysis is reported in Supplemental Table 1-Supplement 2.

**Figure 1 supplement 3.**
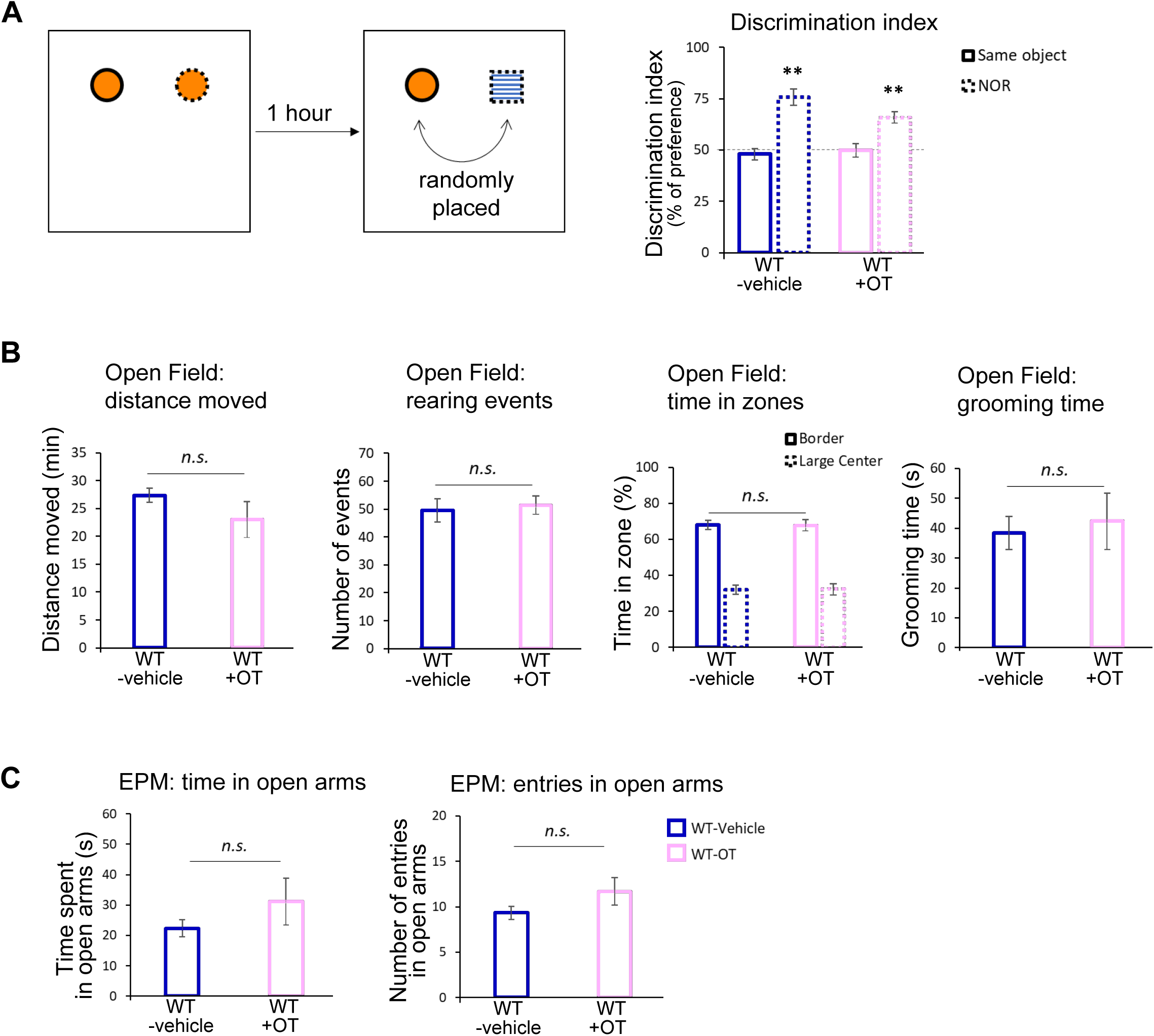
Behavioral tests in male WT mice having been OT-treated or vehicle-treated in neonates. Behavioral analysis of male OT-treated versus vehicle-treated WT mice. (A) Novel object recognition (NOR) test allows to assess the non-social memory. Training simply involves visual exploration of two identical objects, while after one hour the test session involves replacing one of the previously explored objects with a novel object. Because mice have an innate preference for novelty, a mouse that remembers the familiar object (same object) will spend more time exploring the novel object (new object) conducting to a discrimination index significantly different from 50%. A similar discrimination index is observed in vehicle or OT-treated WT males. (B) Open field (OF) test measures locomotor activity and vertical activity (rearing) and anxiety-related behavior (time in zones and grooming): no significant differences have been detected in all those activities between OT-treated and vehicle-treated WT male mice. (C) Elevated Plus Maze (EPM) test allows to measure anxiety (time spent in open arms and number of entries in open arms) and shows a similar behavior with a tendency to spend more time and enter more often in open arms in WT+OT males, suggesting an anxiolytic effect of OT. Data in histograms report mean± SEM. Mann-Whitney test,* P<0.05. Statistical analysis is reported in Supplemental Table S2-3.

**Figure 5-figure supplement 1:**
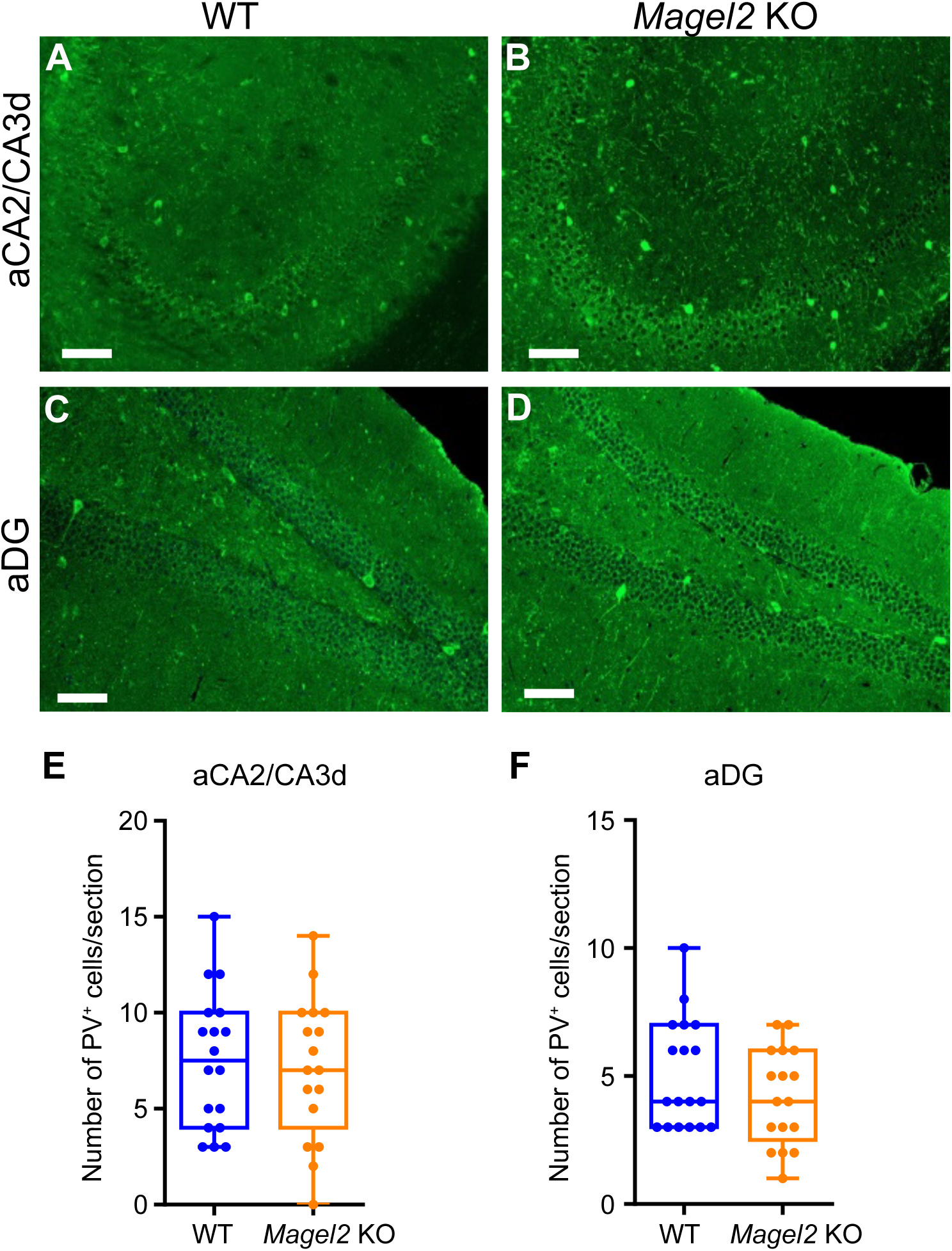
Quantification of paravalbumin (PV) immunopositive cells in the anterior hippocampus region of Magel2 KO adult mice compared with WT mice. (A-D) PV immunolabeling on coronal hippocampal sections at the level of aCA2/CA3d and aDG regions in WT (A-C) and Magel2 KO (B-D). (E-F) Number of PV+ cells by section in both aCA2/CA3d (E) and aDG (F) and comparing WT (N=4) with Magel2 KO (N=4) mice (M-N). Scale bar: 100µm. Data represented in whisker-plots report the number of PV+ cells by sections (8 sections/ hippocampus) with Q2(Q1, Q3) for each genotype and scattered plots showing individual data points. Mann-Whitney Test.

**Figure 6-figure supplement 1.**
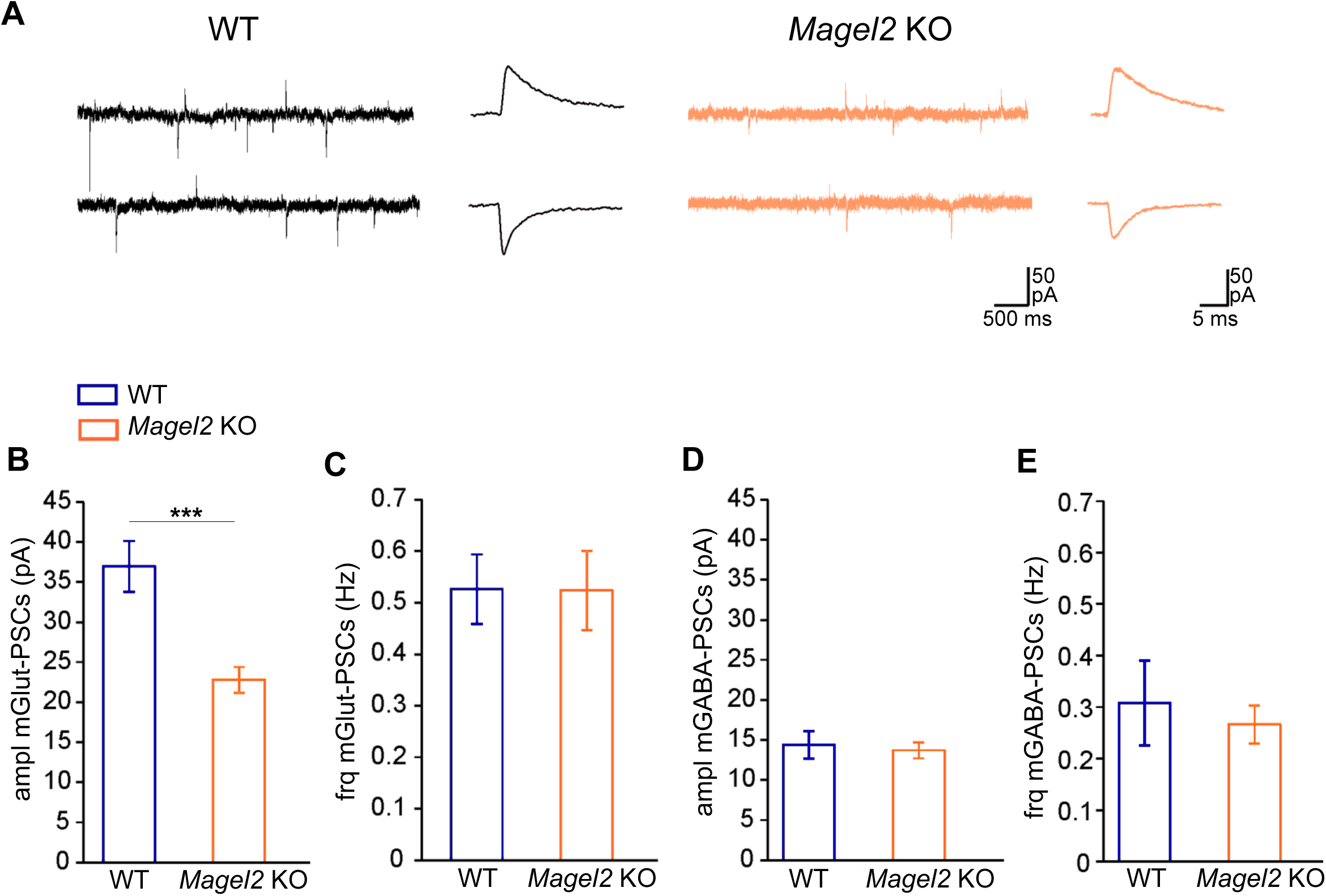
Miniatures Glutamatergic and GABAergic synaptic activitiy of aCA3 pyramidale neurons in the anterior hippocampus region of Magel2 KO juvenile mice compared with WT. (A) Representative whole-cell recordings of miniature Glutamatergic (inward) and GABAergic (outward) currents (holding potential=-45mV) in juvenile WT and Magel2 KO aCA3 pyramidal neurons. (B-E) The amplitudes (B and D) and frequencies (C and E) of the miniatures glutamatergic (B and C) and GABAergic (D and E) postsynaptic currents. GABAergic frequency (Hz) (WT n=16; Magel2 KO n=17), GABAergic amplitude (pA) (WT n=16; Magel2 KO n=17) and glutamatergic frequency (WT n=14; Magel2 KO n=18) are similar but glutamatergic amplitude (pA) is significantly different. WT (N=5, n=14); Magel2 KO (N=5, n=18). N: number of mice and n: number of recorded cells. Data represented in histograms are mean ± SEM. Mann Whitney test, ***P< 0.001. Statistical analysis is reported in Supplemental Table 6-Supplement 1.

**Figure 6-figure supplement 2.**
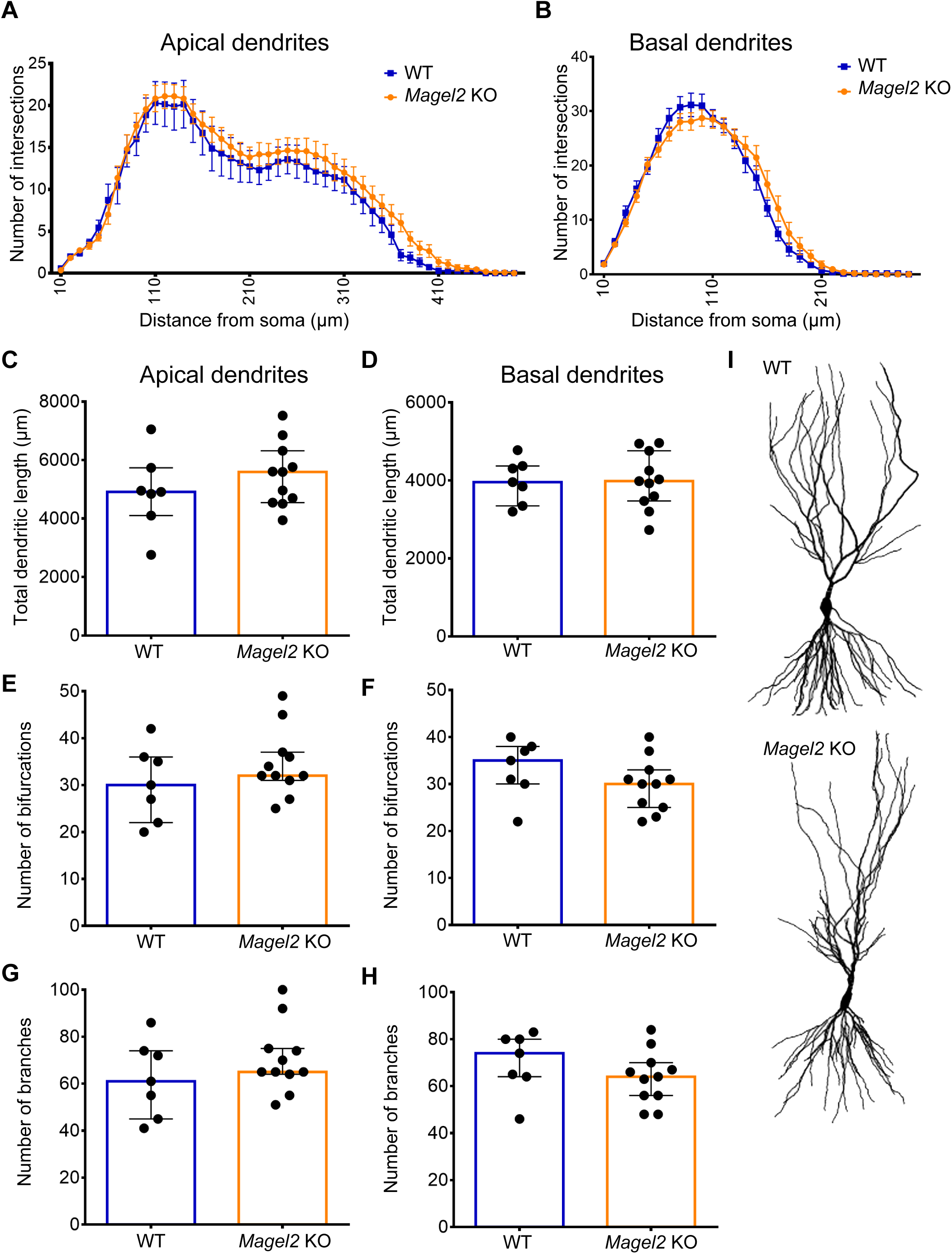
Morphology of aCA3 recorded pyramidal neurons in Magel2 KO versus WT mice. (A) Representative distribution of apical and (B) basal dendritic complexity obtained with Sholl analysis. (C) Quantification of total length of the apical and (D) basal dendrites. (E) Quantification of the mean number of bifurcations in apical and (F) basal dendrites. (G) Quantification of the mean number of total branches in apical and (H) basal dendrites following 3D reconstructions from CA3 pyramidal neurons recorded in previous experiments (Figure 6). (I) Representative WT and Magel2 KO reconstructed CA3 pyramidal neurons. Histograms indicate Q2(Q1, Q3). Each dot represents a neuron (n). WT (N=3, n=7), Magel2 KO (N=3, n=11). N: number of mice and n: number of recorded cells. Mann-Whitney test. Statistical analysis is reported in Supplemental Table 6-Supplement 2.

**Figure 7-Supplement 1.**
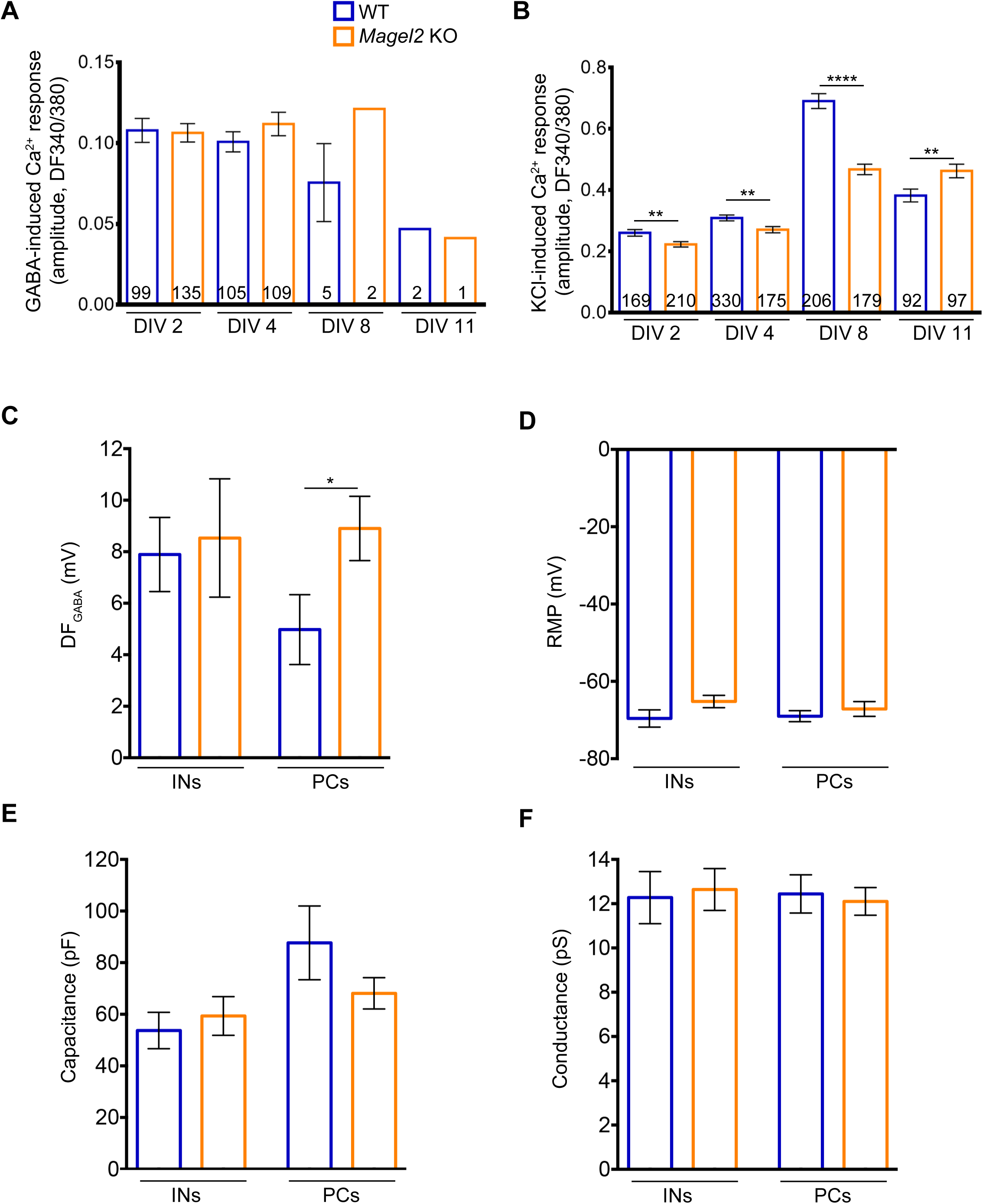
Parameters to validate the in vitro Calcium imaging analysis and the DFGABA study in Magel2 KO versus WT mice. (A) Amplitude of GABA-induced and (B) KCl-induced Ca2+ peaks in WT and Magel2 KO E18 neurons at the different DIVs. Number of responsive cells is reported in the bars. Data are from two to five different preparations, 6-8 embryos each preparation (six to fifteen coverslips analyzed). (C-F) Electrophysiological parameters of pyramidal cells (PCs) and interneurons (INs) recorded at P7 in Magel2 KO versus WT CA3 region of hippocampus. (C) Driving force for GABA (DFGABA) measured with single GABAA channels in cell-attached mode in INs and PCs. (D) Resting membrane potential (RMP) of INs and PCs measured in whole-cell current-clamp mode. (E) Cell capacitance of recorded neurons. (F) Conductance of the GABAA channels recorded in cell-attached mode (presented in Figure 8C). Histograms report mean ± SEM. Unpaired t test with Welch’s correction: *P<0-05, **P<0.01, ****P<0.0,001. Statistical analysis is reported in Supplemental Table 7-Supplement 1.

## SUPPLEMENTAL INFORMATION

### METHODS AND MATERIALS

#### Animals

Magel2 is an imprinted gene, with a monoallelic paternal expression. However, to overcome the phenotypic heterogeneity of the heterozygous +m/-p *Magel2 ^tm1.1Mu^*^s^ mouse, due to the stochastic expression of the maternal *Magel2* allele when the paternal allele is deleted ^67^, *Magel2 ^tm1.1Mus^* homozygous (-/-) mice were used. We made this choice to obtain a greater homogeneity in the values, allowing a better analysis of the effects of the mutation.

*Magel2 ^tm1.1Mus^*+/+ (WT) and *Magel2 ^tm1.1Mus^*-/- (*Magel2-KO*) mice were stabulated in standard conditions, with *ad-libitum* access to food and water. Mice were handled and cared for in accordance with the Guide for the Care and Use of Laboratory Animals (N.R.C., 1996) and the European Communities Council Directive of September 22th 2010 (2010/63/EU, 74). All efforts were made to minimize the number of animals used. *Magel2*-deficient mice were generated as previously described ^16^. Due to the parental imprinting of *Magel2* (paternally expressed only), to obtain heterozygote mice (+m/-p), males carrying the mutation on the maternal allele (-m/+p) were crossed with wild-type C57BL/6J females.

To obtain homozygote mice, *Magel2-KO* homozygote males and females were crossed. Importantly, we checked that *Magel2-KO* mothers had a similar maternal behavior as WT mothers. All mice were genotyped by PCR starting from DNA extracted from tail snips (around 3 mm), using the following couples of primers: Ml2KO F (5’-CCCTGGGTTGACTGACTCAT-3’) and Ml2KO R (5’-TCTTCTTCCTGGTGGCTTTG-3’) to discriminate the mutant allele from the WT, 71456 F (5’-CACTCGATCACGTATGGCTCCATCA-3’) and 71457 R (5’-GATGGCAGGCACTGACTTACATGCTG-3’) to discriminate the heterozygous from the homozygous mice.

#### Behavior

All the behavioral tests were performed by Phenotype Expertise, Inc. (France). For all tests, animals were first acclimated to the behavioral room for 30 minutes.

##### Elevated-Plus Maze

The EPM is used to assess anxiety state of animals. The device consists of a labyrinth of 4 arms 5 cm wide located 80 cm above the ground. Two opposite arms are open (without wall) while the other two arms are closed by side walls. The light intensity was adjusted to 20 Lux on the open arms. Mice were initially placed on the central platform and left free to explore the cross-shaped labyrinth for 5 minutes. Maze was cleaned and wiped with H_2_O and with 70% ethanol between each mouse. Animal movement was video-tracked using Ethovision software 11.5 (Noldus). Time spent in open and closed arms, the number of entries in open arms, as well as the distance covered, are directly measured by the software.

##### Open-field

Open-field test was performed in a 40 x 40 cm square arena with an indirect illumination of 60 lux. Mouse movement was video-tracked using Ethovision software 11.5 (Noldus) for 10 minutes. Total distance traveled and time in center (exclusion of a 5 cm border arena) are directly measured by the software. Grooming (time and events) and rearing were manually counted in live using manual functions of the software, by an experimented behaviorist. The open-field arena was cleaned and wiped with H_2_O and with 70% ethanol between each mouse.

##### New object recognition

The arena used for the novel object recognition test was the same used for the open-field test. The arena was cleaned and wiped with 70% ethanol between each mouse. Two identical objects (50 ml orange corning tube) were placed in the opposite corners of the arena, 10 cm from the side walls. The tested mouse was placed at the opposite side of the arena and allowed to explore the arena for 10 min. After 1h, one object was randomly replaced with another novel object, which was of similar size but differ in the shape and color with the previous object (white and blue lego bricks). Then, the same mouse was placed in the arena and allowed to explore the two objects (a new and an “old” familiar object) for 10 min. The movement of the mice was video-tracked with Ethovision 11.5 software. Time of exploration of both objects (nose located in a 2 cm area around object) was automatically measured by the software.

##### Three-chamber social preference test

The test was performed as described previously ^68^. The three-chamber apparatus consisted of a Plexiglas box (50×25 cm) with removable floor and partitions dividing the box into three chambers with 5-cm openings between chambers. The task was carried out in four trials. The three-chambers apparatus was cleaned and wiped with 70% ethanol between each trial and each mouse.

In the first trial (habituation), a test mouse was placed in the center of the three-chamber unit, where two empty wire cages were placed in the left and right chambers to habituate the test mouse to arena. The mouse was allowed to freely explore each chamber. The mouse was video-tracked for 5 min using Ethovision software. At the end of the trial, the animal was gently directed to the central chamber with doors closed. In the second trial (social exploration), a 8-weeks old C57BL/6J mouse (S1) was placed randomly in one of the two wire cages to avoid a place preference. The second wire cage remained empty (E). Then, doors between chambers were opened and the test mouse was allowed to freely explore the arena for 10 min. At the end of the trial, animal was gently directed to the central chamber with doors closed. A second 8-weeks old C57BL/6J mouse (S2) was placed in the second wire cage for the third trial (social discrimination). Thus, the tested mouse had the choice between a familiar mouse (S1) and a new stranger mouse (S2) for 10 min. At the end of the trial, the mouse was returned to home-cage for 30 min. For the fourth trial (short-term social memory), S2 was replaced by a new stranger mouse (S3), the familiar mouse (S1) staying the same. Then tested mouse was allowed to freely explore the arena for 10 min. Time spent in each chamber and time of contact with each wire cage (with a mouse or empty) were calculated using Ethovision software. The measure of the real social contact is represented by the time spent in nose-to-nose interactions with the unfamiliar or familiar mouse. This test was performed using grouped-house mice of 4 months old.

#### Primary hippocampal cultures

Embryonic day 18 dissociated hippocampal neurons were obtained from wild-type and *Magel2-KO* timed pregnant mice as previously described ^69^ with slightly modifications here described. Briefly, the hippocampi of E18 embryos were dissociated by an enzymatic treatment (0.25% trypsin for 18 min at 37°C) followed by mechanic dissociation with a fire-smoothed Pasteur pipette or p1000µl/p200µl tips. For calcium imaging experiments, 200 000 cell/well (in MW 6 wells) were plated on round 26 mm glass coverslips pre-coated with poly-L-lysine containing Neurobasal medium (Life Technologies) augmented with B27 supplement (2% v/v; Life Technologies), L-glutamine (2mM), penicillin/streptomycin (100U/ml) and 25µM Glutamate. This media was replaced with glutamate-free media after 5 hours. Neurons were then maintained at 37°C in humidified atmosphere (95% air and 5% CO_2_), and half of the medium was refreshed twice a week.

#### Calcium imaging recordings

For calcium imaging experiments, hippocampal neurons were loaded with the membrane-permeable fluorescent Ca^2+^ indicator Fura-2/AM (1 μM; SigmaAldrich) for 40 min at 37°C, 5% CO_2_. The cells were then placed into the recording chamber of an inverted microscope (Axiovert 100, Zeiss), washed with the extracellular recording solution, KRH buffer, and imaged through a 40x objective (Zeiss). Fura-2/AM was excited at 380 nm and at 340 nm through a Polychrom V, (TILL Photonics GmbH) controlled by the TillVisION software 4.01. Emitted light was acquired at 505nm at 1Hz, and images collected with a CCD Imago-QE camera (TILL Photonics GmbH). The fluorescence ratio F340/380 (ΔF340/380) was used to express Ca^2+^ concentrations in regions of interest (ROI) corresponding to neuronal cell bodies. 100µM GABA was administered in the recording solution and temporal changes in ΔF340/380 were followed. Increases in ΔF340/380 higher than 0.04 units were considered reliable Ca^2+^ responses. After wash with KRH buffer and recover, KCl (50mM) was administered to identify viable neurons. Responses with a ΔF340/380 smaller than 0.1 units were excluded from the analyses. From DIV8 on, 1 μM TTX (Tocris, cat #1069) was added to this extracellular recording solution.

#### Hippocampal slice preparation and electrophysiological recordings

Brains were removed and immersed into ice-cold (2-4°C) artificial cerebrospinal fluid (ACSF) with the following composition (in mM): 126 NaCl, 3.5 KCl, 2 CaCl_2_, 1.3 MgCl_2_, 1.2 NaH_2_PO_4_, 25 NaHCO_3_ and 11 glucose, pH 7.4 equilibrated with 95% O_2_ and 5% CO_2_. Hippocampal slices (400 µm thick) were cut with a vibrating microtome (Leica VT 1000s, Germany) in ice cold oxygenated choline-replaced ACSF and were allowed to recover at least 90 min in ACSF at room (25°C) temperature. Slices were then transferred to a submerged recording chamber perfused with oxygenated (95% O_2_ and 5% CO_2_) ACSF (3 ml/min) at 34°C.

##### Whole-cell patch clamp recordings

were performed from P20-P25 CA3 pyramidal neurons in voltage-clamp mode using an Axopatch 200B (Axon Instrument, USA). To record the spontaneous and miniature synaptic activity, the glass recording electrodes (4-7 MΩ) were filled with a solution containing (in mM): 100 KGluconate, 13 KCl, 10 HEPES, 1.1 EGTA, 0.1 CaCl_2_, 4 MgATP and 0.3 NaGTP. The pH of the intracellular solution was adjusted to 7.2 and the osmolality to 280 mOsmol l^-1^. The access resistance ranged between 15 to 30 MΩ. With this solution, the GABA_A_ receptor-mediated postsynaptic current (GABAA-PSCs) reversed at −70mV. GABA-PSCs and glutamate mediated synaptic current (Glut-PSCs) were recorded at a holding potential of −45mV. At this potential GABA-PSC are outwards and Glut-PSCs are inwards. Spontaneous synaptic activity was recorded in control ACSF and miniature synaptic activity was recorded in ACSF supplemented with tetrodotoxin (TTX, 1µM). Spontaneous and miniature GABA-PSCs and Glut-PSCs were recorded with Axoscope software version 8.1 (Axon Instruments) and analyzed offline with Mini Analysis Program version 6.0 (Synaptosoft).

##### Single GABA_A_ channel recordings

were performed at P1, P7 and P15 visually identified hippocampal CA3 pyramidal cells in cell-attached configuration using Axopatch-200A amplifier and pCLAMP acquisition software (Axon Instruments, Union City, CA). Data were low-pass filtered at 2 kHz and acquired at 10 kHz. The glass recording electrodes (4-7 MΩ) were filled with a solution containing (in mM) for recordings of single GABA_A_ channels: NaCl 120, KCl 5, TEA-Cl 20, 4-aminopyridine 5, CaCl_2_ 0.1, MgCl_2_ 10, glucose 10, Hepes-NaOH 10. The pH of pipette solutions was adjusted to 7.2 and the osmolality to 280 mOsmol l^-1^. Analysis of currents trough single channels and current-voltage relationships were performed using Clampfit 9.2 (Axon Instruments) as described by ^70^.

#### Morphological analysis

During electrophysiological recordings, biocytin (0.5%, Sigma, USA) was added to the pipette solution for post hoc reconstruction. Images were acquired using a Leica SP5 X confocal microscope, with a 40x objective and 0,5 µm z-step. Neurons were reconstructed tree-dimensionally using Neurolucida software version 10 (MBF Bioscience) from 3D stack images. The digital reconstructions were analyzed with the software L-Measure to measure the number of primary branches and the total number of ramifications of each neuron ^71^. Comparisons between groups were done directly in L-Measure.

#### Immunohistochemistry and quantification

WT and mutant mice were deeply anaesthetized with intraperitoneal injection of the ketamine/xylazine mixture and transcardially perfused with 0.9% NaCl saline followed by Antigenfix (Diapath, cat #P0014). Brains were post-fixed in Antigenfix overnight at 4°C and included in agar 4%. 50 μm-thick coronal sections were sliced using a vibratom (Zeiss) and stored in PBS at 4°C. Floating slices (of the hippocampal region corresponding to slices 68 to 78 on Allan Brain Atlas) were incubated for 1 hour with blocking solution containing 0.1% (v/v) Triton X-100, 10% (v/v) normal goat serum (NGS) in PBS, at room temperature. Sections were then incubated with primary antibodies diluted in incubation solution (0.1% (v/v) Triton X-100, 3% (v/v) NGS, in PBS), overnight at 4C°. After 3 x 10 min washes in PBS, brain sections were incubated with secondary antibodies diluted in the incubation solution, for 2 hours at RT. Sections were washed 3 x 10 min in PBS and mounted in Flouromount-G (EMS, cat #17984-25). Primary antibodies used were: rabbit polyclonal anti-cFos (1:5000, Santa Cruz Biotech, cat #ab190289), goat polyclonal anti-Sst (D20) (1:500, Santa Cruz Biotech, cat #sc-7819), mouse monoclonal anti-Sst (H-11) (1:500, Santa Cruz Biotech, cat #sc-74556), goat polyclonal anti-PV (1:6000, SWANT, cat #PVG213). Fluorochrome-conjugated secondary antibodies used were: goat anti-rabbit Alexa Fluor 647 (1:500, Invitrogen, cat # A32733), goat anti-rabbit Alexa Fluor 488 (1:500, Invitrogen, cat #A-31565), goat anti-mouse Alexa Fluor 488 (1:500, Invitrogen, cat #A21121), donkey anti-goat Alexa Fluor 488 (1:500, Invitrogen, cat # A32814).

For c-Fos, PV and SST quantification, images were acquired using a fluorescence microscope (Zeiss Axioplan 2 microscope with an Apotome module), and z stacks of 8 µm were performed for each section. Counting wsa performed on the right and left hippocampus for 5-7 sections (cFos) or 7-9 sections (PV, SST) per animal in the hippocampal regions indicated on the figures and corresponding to slices 68 to 78 on Allen Brain Atlas.

#### OT binding assay

Briefly, slides were pre-incubated for 5 minutes in a solution of 0.2% paraformaldehyde in phosphate-buffered saline (pH 7.4), and rinsed twice in 50 mM Tris HCl + 0.1% BSA buffer. Slides were then put in a humid chamber and covered with 400 µL of incubation medium (50 mM Tris HCl, 0.025% bacitracin, 5 mM MgCl2, 0,1% BSA) containing the radiolabeled I [125] OVTA (Perkin Elmer), at a concentration of 10 pM. After a 2h incubation under gentle agitation, the incubation medium is removed and slides are rinsed twice in ice-cold incubation medium and a third time in ice cold distilled water. Each slide is then dried in a stream of cool air, and placed in an X-ray cassette in contact with a KODAK film for 3 days.

ROIs were chosen and analyzed through ImageJ, using Paxinos’ Mouse Brain Atlas as a reference to find the brain areas of interest. To remove background noise caused by nonspecific binding, each slide was compared with its contiguous one, which had been incubated in presence of an excess of “cold” oxytocin (2 μM). Net grey intensity was quantified and then converted to nCi/mg tissue equivalent using a calibration curve. For each region, a minimum of 4 slices per brain were included in the analysis. Data plotted on graphs are the differences between the total and the nonspecific binding. Right and left hemispheres were kept separate.

#### Chromogenic In situ Hybridization

The two probes used are synthetic oligonucleotide probes complementary to the nucleotide sequence 1198 – 2221 of Oxtr (NM_001081147.1) (Oxtr-E4-C2, ACD Cat #411101-C2) and 3229 – 4220 of Magel2 (NM_013779.2) (Magel2-01, ACD Cat #535901). Briefly, slides were fixed in 4% paraformaldehyde in PBS (pH 9.5) on ice for 2 hours and dehydrated in increasing concentrations of alcohol, then stored in 100% ethanol overnight at −20°C. The slides were air dried for 10 minutes, then pretreated in target retrieval solution (ref. 322001, ACD) for 5 minutes while boiling, after which, slides were rinsed 2 times in water followed by 100% ethanol and then air dried. A hydrophobic barrier pen (ImmEdge) was used to create a barrier around selected sections. Selected sections were then incubated with protease plus (ref. 322331, ACD) for 15 minutes in a HybEZ oven (ACD) at 40°C, followed by water washes. The sections were then hybridized with the probe mixture at 40°C for 2 hr per slide. Unbound hybridization probes were removed by washing 2 times in wash buffer. After hybridization, sections were subjected to signal amplification using the HD 2.5 detection Kit following the kit protocol. Hybridization signal was detected using a mixture of fast-RED solutions A and B (60:1) for Oxtr-E4-C2 and a mixture of Fast-GREEN solutions A and B (50:1) for Magel2-01. The slides were then counterstained with Gill’s hematoxylin and air-dried in a 60°C oven for 15 min. Slides were cooled and cover-slipped with Vectamount TM (Vector Laboratories, Inc. Burlingame, CA). Slides were imaged at 4x and 20x on a bright field microscope (Keyence BZ-X710, Keyence Corp., Osaka, Japan). Hippocampal sections were investigated for colocalization of Oxtr (red) with Magel2 (blue-green) transcripts.

#### Western Blot

P7 mice were sacrificed and hippocampi were dissected and rapidly frozen in liquid nitrogen and stored at −80°C until protein extraction. Hippocampi were lysed in lysis buffer (50 mM Tris/HCl, pH 7.5, 1 mM EGTA, 1 mM EDTA, 50 mM sodium fluoride, 5 mM sodium pyrophosphate, 1 mM sodium orthovanadate, 1% (w/v) Triton-100, 0.27 M sucrose, 0.1% (v/v) 2-mercaptoethanol, and protease inhibitors (complete protease inhibitor cocktail tablets, Roche, 1 tablet per 50 mL)) and protein concentrations were determined following centrifugation of the lysate at 16,000 x g at 4 °C for 20 minutes using the Bradford method with bovine serum albumin as the standard. Tissue lysates (15 µg) in SDS sample buffer (1X NuPAGE LDS sample buffer (Invitrogen), containing 1% (v/v) 2-mercaptoethanol) were subjected to electrophoresis on polyacrylamide gels and transferred to nitrocellulose membranes. The membranes were incubated for 30 min with TBS-Tween buffer (TTBS, Tris/HCl, pH 7.5, 0.15 M NaCl and 0.2% (v/v) Tween-20) containing 5% (w/v) skim milk. The membranes were then immunoblotted in 5% (w/v) skim milk in TTBS with the indicated primary antibodies overnight at 4°C. The blots were then washed six times with TTBS and incubated for 1 hour at room temperature with secondary HRP-conjugated antibodies diluted 5000-fold in 5% (w/v) skim milk in TTBS. After repeating the washing steps, the signal was detected with the enhanced chemiluminescence reagent. Immunoblots were developed using ChemiDoc™ Imaging Systems (Bio-Rad). Primary antibodies used were: anti-KCC2 phospho-Ser940 (Thermo Fisher Scientific, cat #PA5-95678), anti-KCC2 phospho-Thr1007 (Thermo Fisher Scientific, cat #PA5-95677), anti-Pan-KCC2, residues 932-1043 of human KCC2 (NeuroMab, cat #73-013), anti(neuronal)-β-Tubulin III (Sigma-Aldrich, cat #T8578). Horseradish peroxidase-coupled (HRP) secondary antibodies used for immunoblotting were from Pierce. Figures were generated using Photoshop and Illustrator (Adobe). The relative intensities of immunoblot bands were determined by densitometry with ImageJ software.

### Statistical Analysis

Statistical analyses were performed using GraphPad Prism (GraphPad Software, Prism 7.0 software, Inc, La Jolla, CA, USA). All statistical tests were two-tailed and the level of significance was set at P<0.05. Appropriate tests were conducted depending on the experiment; tests are indicated in the figure legends or detailed in supplementary statistical file. Values are indicated as Q2 (Q1, Q3), where Q2 is the median, Q1 is the first quartile and Q3 is the third quartile when non-parametric tests were performed and scatter dot plots report Q2 (Q1,Q3) or as mean ± SEM when parametric tests were performed usually in histograms. N refers to the number of animals or primary culture preparations, while n refers to the number of brain sections or hippocampi or cells recorded.

Mann-Whitney (MW) non-parametric test or t-test (parametric test) were performed to compare two matched or unmatched groups. ANOVA or Kruskal-Wallis tests were performed when the different groups have been experienced in the same experimental design only; if this was not the case, MW or t-test were used. One-way ANOVA followed by Bonferroni or Dunnett’s or Tukey’s post-hoc tests were used to compare three or more independent groups. Two-way ANOVA followed by Bonferroni post-hoc test was performed to compare the effect of two factors on unmatched groups. *: p< 0.05; **: p <0.01; ***: p<0.001; ****: p<0.0001. All the statistical analyses (corresponding to each figure) are reported in a specific file.

---

**Table 1.**

| Figure | Parameter | Test specifications |  | Genotype | Treatment | Sex | Number of individuals | Mean ± SEM | Statistical test | p-value | Median (Q1,Q3) | Statistical test | p-value |
| --- | --- | --- | --- | --- | --- | --- | --- | --- | --- | --- | --- | --- | --- |
| 1B | Interaction time (sniffing in seconds) | Social exploration | S1 | WT | - | male | 12 | 78.0 ± 7.7 | paired t-test | p=0.00134 | 73.48 (52.34, 100.58) | Wilcoxon Signed Rank Test | p<0.001 |
|  |  |  | Empty |  |  |  |  | 34.5 ± 6.6 |  |  | 27.52 (16.72, 56.22) |  |  |
|  |  | Social discrimination | S1 | WT | - | male |  | 33.7 ± 4.7 |  | p=0.000129 | 28.04 (21.00, 45.96) |  | p<0.001 |
|  |  |  | S2 |  |  |  |  | 70.7 ± 7.4 |  |  | 78.68 (53.50, 89.12) |  |  |
|  |  | Short-term social memory | S1 | WT | - | male |  | 28.0 ± 6.0 |  | p=0.0263 | 21.08 (14.34, 33.84) |  | p=0.042 |
|  |  |  | S3 |  |  |  |  | 77.2 ± 16.3 |  |  | 60.28 (32.78, 109.96) |  |  |
|  |  | Social exploration | S1 | Magel2 KO | - | male | 9 | 68.1 ± 6.8 |  | p=0.000133 | 65.04 (51.80, 81.44) |  | p=0.004 |
|  |  |  | Empty |  |  |  |  | 31.5 ± 5.0 |  |  | 24.64 (21.32, 44.88) |  |  |
|  |  | Social discrimination | S1 | Magel2 KO | - | male |  | 30.0 ± 4.3 |  | p=0.0169 | 25.92 (21.76, 43.56) |  | p=0.012 |
|  |  |  | S2 |  |  |  |  | 61.0 ± 7.6 |  |  | 65.92 (44.48, 71.60) |  |  |
|  |  | Short-term social memory | S1 | Magel2 KO | - | male |  | 38.2 ± 6.4 |  | p=0.472 | 37.68 (22.88, 47.20) |  | p=0.359 |
|  |  |  | S3 |  |  |  |  | 43.9 ± 7.1 |  |  | 46.16 (22.20, 55.56) |  |  |
| 1C | Interaction time (sniffing in seconds) | Social exploration | S1 | WT | vehicle | male | 18 | 83.5 ± 6.5 | paired t-test | p=0.00179 | 79.48 (70.24, 95.40) | Wilcoxon Signed Rank Test | p=0.001 |
|  |  |  | Empty |  |  |  |  | 49.2 ± 5.6 |  |  | 41.88 (33.00, 65.74) |  |  |
|  |  | Social discrimination | S1 | WT | vehicle | male |  | 44.2 ± 3.8 |  | p=0.000291 | 45.96 (28.34, 53.86) |  | p<0.001 |
|  |  |  | S2 |  |  |  |  | 84.5 ± 6.9 |  |  | 79.16 (58.58, 107.38) |  |  |
|  |  | Short-term social memory | S1 | WT | vehicle | male |  | 34.0 ± 3.3 |  | p=0.000571 | 31.60 (25.06, 39.00) |  | p=0.001 |
|  |  |  | S3 |  |  |  |  | 67.1 ± 6.4 |  |  | 67.44 (41.44, 97.10) |  |  |
|  |  | Social exploration | S1 | WT | OT | male | 10 | 88.3 ± 6.3 |  | p<0.0001 | 83.68 (70.94, 98.44) |  | p=0.002 |
|  |  |  | Empty |  |  |  |  | 42.0 ± 2.3 |  |  | 44.76 (33.98, 48.36) |  |  |
|  |  | Social discrimination | S1 | WT | OT | male |  | 49.9 ± 4.9 |  | p=0.00776 | 49.64 (36.72, 63.86) |  | p=0.010 |
|  |  |  | S2 |  |  |  |  | 84.2 ± 10.5 |  |  | 71.84 (57.14, 103.48) |  |  |
|  |  | Short-term social memory | S1 | WT | OT | male |  | 36.4 ± 3.4 |  | p=0.00917 | 35.08 (27.90, 47.72) |  | p=0.020 |
|  |  |  | S3 |  |  |  |  | 68.8 ± 7.4 |  |  | 71.84 (46.10, 86.92) |  |  |
| 1D | Interaction time (sniffing in seconds) | Social exploration | S1 | Magel2 KO | vehicle | male | 19 | 65.8 ± 5.1 | paired t-test | p<0.0001 | 69.44 (58.72, 79.04) | Wilcoxon Signed Rank Test | p<0.001 |
|  |  |  | Empty |  |  |  |  | 31.0 ± 3.2 |  |  | 26.88 (22.80, 40.40) |  |  |
|  |  | Social discrimination | S1 | Magel2 KO | vehicle | male |  | 29.5 ± 4.3 |  | p<0.0001 | 26.80 (14.64, 36.96) |  | p<0.001 |
|  |  |  | S2 |  |  |  |  | 73.5 ± 7.0 |  |  | 74.64 (43.12, 92.64) |  |  |
|  |  | Short-term social memory | S1 | Magel2 KO | vehicle | male |  | 38.8 ± 4.0 |  | p=0.259 | 42.08 (25.36, 52.08) |  | p=0.374 |
|  |  |  | S3 |  |  |  |  | 44.2 ± 3.1 |  |  | 43.20 (37.52, 50.64) |  |  |
|  |  | Social exploration | S1 | Magel2 KO | OT | male | 19 | 89.1 ± 6.0 |  | p<0.0001 | 84.40 (68.24, 113.60) |  | p<0.001 |
|  |  |  | Empty |  |  |  |  | 36.2 ± 4.5 |  |  | 34.56 (21.12, 50.96) |  |  |
|  |  | Social discrimination | S1 | Magel2 KO | OT | male |  | 34.0 ± 4.7 |  | p<0.0001 | 33.52 (17.44, 43.60) |  | p<0.001 |
|  |  |  | S2 |  |  |  |  | 87.2 ± 6.2 |  |  | 86.16 (75.12, 98.00) |  |  |
|  |  | Short-term social memory | S1 | Magel2 KO | OT | male |  | 32.4 ± 3.8 |  | p=0.000702 | 28.96 (22.40, 38.40) |  | p<0.001 |
|  |  |  | S3 |  |  |  |  | 58.6 ± 6.5 |  |  | 53.32 (43.76, 64.80) |  |  |

**Table 1-Supplement 1, 2,3.**
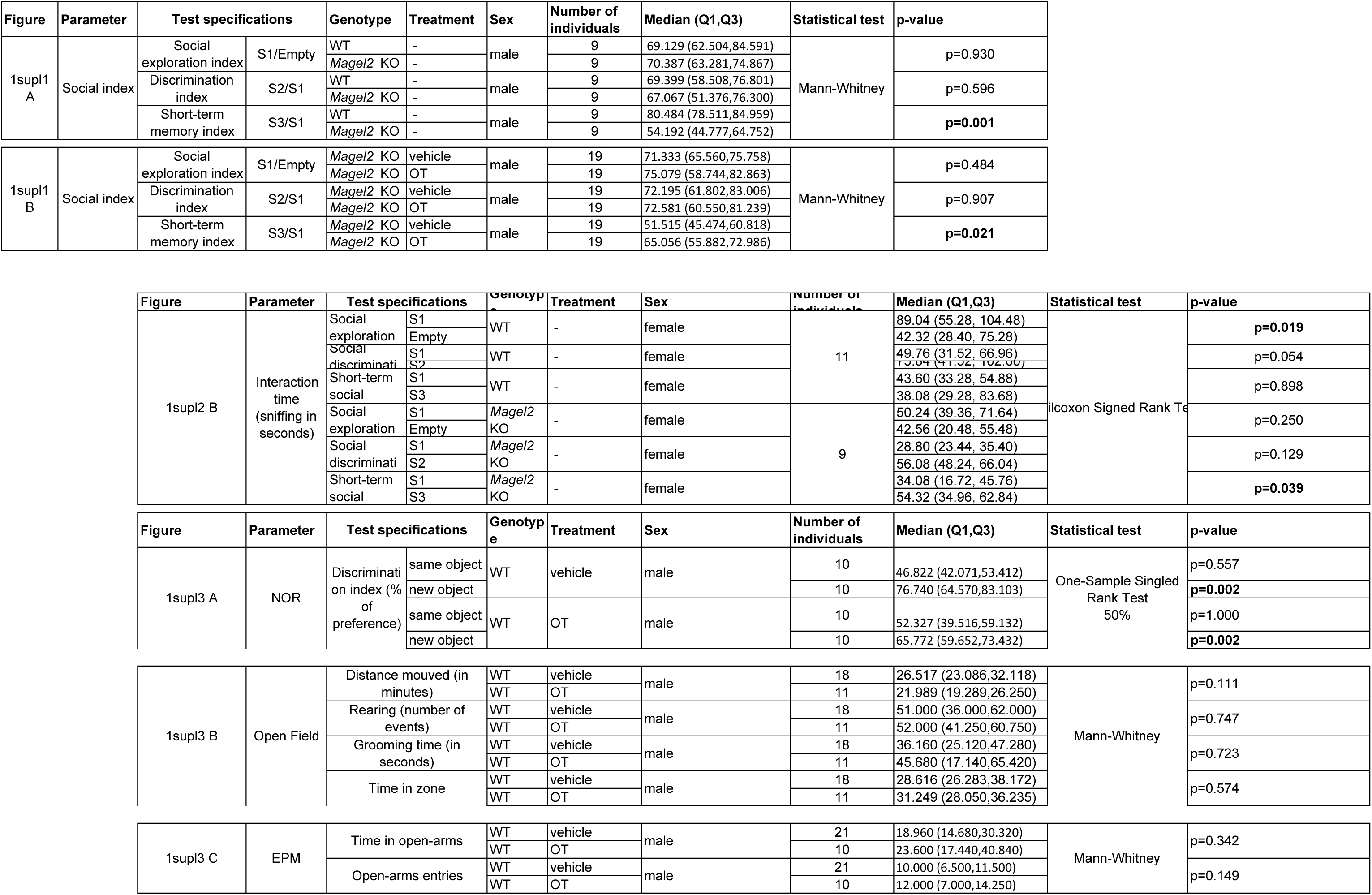

**Table 2.**

| Figure | Parameter | Brain region | Genotype | Behavioral test | Treatment | Gender | Number of individuals | Number of sections | Median (Q1,Q3) | Statistical test | p-value |
| --- | --- | --- | --- | --- | --- | --- | --- | --- | --- | --- | --- |
| 3D | Number of cFos-positive cells | aCA2/CA3d | WT | -SI | - | male | 3 | 18 | 24.5 (19.5,28) | one-way ANOVA,<br>Dunnett's post-hoc test |  |
|  |  |  | WT | +SI | - | male | 4 | 24 | 44.5 (34,56) |  | **WT-SI vs WT +SI, p<0.0001** |
|  |  |  | *Magel2* KO | -SI |  | male | 4 | 20 | 25 (22.2,28) |  | **KO-SI vs KO +SI, p<0.0001** |
|  |  |  | *Magel2* KO | +SI | - | male | 4 | 24 | 55 (48,75) |  | **WT+SI vs *Magel2* KO+SI, p=0.03** |
| 3E | Number of cFos-positive cells | aDG | WT | -SI | - | male | 3 | 18 | 39.5 (31.7,42) | one-way ANOVA,<br>Dunnett's post-hoc test |  |
|  |  |  | WT | +SI | - | male | 4 | 24 | 63 (53.5,79.5) |  | **WT-SI vs WT+SI, p<0.0001** |
|  |  |  | *Magel2* KO | -SI |  | male | 4 | 20 | 39 (38,46) |  | **KO-SI vs KO +SI, p<0.0001** |
|  |  |  | *Magel2* KO | +SI | - | male | 4 | 24 | 72.5 (58.7,85.2) |  | WT+SI vs *Magel2* KO+SI, n.s. |

**Table 4.**

| Figure | Parameter | Brain region | Genotype | Treatment | Gender | Number of individuals | Number of sections | Median (Q1,Q3) | Statistical test | p-value |
| --- | --- | --- | --- | --- | --- | --- | --- | --- | --- | --- |
| 4B | nCi/mg of tissue equivalent | aCA2/CA3 | WT | vehicle | male | 3 | 24 | 0.07469 (0.06504,0.08431) | one-way ANOVA, Bonferroni's post-hoc test | **WT-vehicle vs *Magel2* KO-vehicle, p<0.0001** |
|  |  |  | *Magel2* KO | vehicle | male | 3 | 24 | 0.1395 (0.1231,0.1593) |  | **WT-vehicle vs *Magel2* KO+OT, p=0.0021** |
|  |  |  | *Magel2* KO | OT | male | 3 | 24 | 0.1254 (0.1057,0.1354) |  | *Magel2* KO-vehicle vs *Magel2* KO+OT, p=0.2232 |
| 4D | nCi/mg of tissue equivalent | vCA1/CA2/CA3 | WT | vehicle | male | 3 | 24 | 0.07877 (0.06478,0.09454) | one-way ANOVA, Bonferroni's post-hoc test | **WT-vehicle vs *Magel2* KO-vehicle, p<0.0001** |
|  |  |  | *Magel2* KO | vehicle | male | 3 | 24 | 0.1378 (0.1284,0.1535) |  | WT-vehicle vs *Magel2* KO+OT, p>0.9999 |
|  |  |  | *Magel2* KO | OT | male | 3 | 24 | 0.08789 (0.06257,0.0942) |  | ***Magel2* KO-vehicle vs *Magel2* KO+OT, p<0.0001** |
| 4F | nCi/mg of tissue equivalent | aDG | WT | vehicle | male | 3 | 24 | 0.1610 (0.1440,0.1953) | one-way ANOVA, Bonferroni's post-hoc test | WT-vehicle vs *Magel2* KO-vehicle, p=0.0722 |
|  |  |  | *Magel2* KO | vehicle | male | 3 | 24 | 0.1274 (0.1196,0.1415) |  | WT-vehicle vs *Magel2* KO+OT, p=0.0590 |
|  |  |  | *Magel2* KO | OT | male | 3 | 24 | 0.1252 (0.1066,0.1517) |  | *Magel2* KO-vehicle vs *Magel2* KO+OT, p>0.9999 |

**Table 5.**
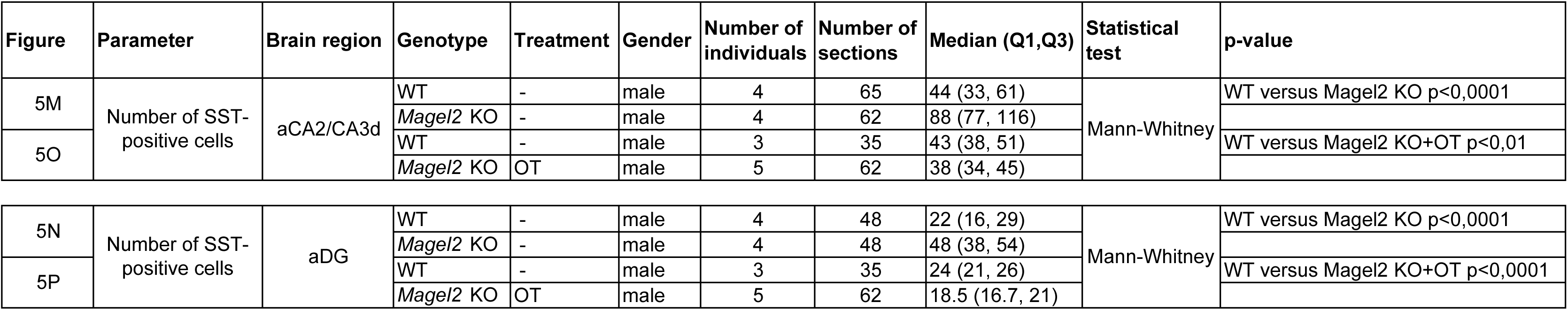

**Table 6.**

| Figure | Parameter | Genotype | Treatment | Gender | Number of individuals | Number of neurons | Median (Q1,Q3) | Statistical test | Comparison | P-value |
| --- | --- | --- | --- | --- | --- | --- | --- | --- | --- | --- |
| 6C | Glut frequency (Hz) | WT | - | male | 7 | 15 | 26 (18, 35) | One way ANOVA + Tukey post-hoc test | WT vs *Magel2* KO | n.s. |
|  |  | *Magel2* KO | - | male | 7 | 18 | 32 (27, 37.5) |  | *Magel2* KO vs KO+OT | p<0.01 |
|  |  | WT | OT | male | 4 | 15 | 8.5 (2.7, 16.2) |  | WT vs WT+OT | p<0.01 |
|  |  | *Magel2* KO | OT | male | 5 | 21 | 11 (17.6, 23.2) |  | WT+OT vs KO+OT | p<0.01 |
| 6E | GABA frequency (Hz) | WT | - | male | 7 | 15 | 13 (5.2, 21) | One way ANOVA + Tukey post-hoc test | WT vs *Magel2* KO | p<0.01 |
|  |  | *Magel2* KO | - | male | 7 | 18 | 23.5 (21.7, 29.2) |  | *Magel2* KO vs KO+OT | p<0.01 |
|  |  | WT | OT | male | 4 | 14 | 11.5 (8, 16.2) |  | WT vs WT+OT | p<0.01 |
|  |  | *Magel2* KO | OT | male | 5 | 21 | 12.5 (10.5, 17.2) |  | WT+OT vs KO+OT | n.s. |
| 6D | Glut amplitude (pA) | WT | - | male | 7 | 15 | 42(18, 61) | One way ANOVA + Tukey post-hoc test | WT vs *Magel2* KO | p<0.05 |
|  |  | *Magel2* KO | - | male | 7 | 16 | 24 (22, 34) |  | *Magel2* KO vs KO+OT NS |  |
|  |  | WT | OT | male | 4 | 15 | 25.5 (21.7, 34) |  | WT vs WT+OT | p<0.01 |
|  |  | *Magel2* KO | OT | male | 5 | 20 | 27.3 (19.3, 32) |  | WT+OT vs KO+OT | p<0.01 |
| 6F | GABA amplitude (pA) | WT | - | male | 7 | 15 | 29 (23.6, 33) | One way ANOVA + Tukey post-hoc test | WT vs *Magel2* KO | n.s. |
|  |  | *Magel2* KO | - | male | 7 | 16 | 24.5 (17.5, 37.5) |  | *Magel2* KO vs KO+OT | n.s. |
|  |  | WT | OT | male | 4 | 15 | 25 (17.4, 30.4) |  | WT vs WT+OT | n.s. |
|  |  | *Magel2* KO | OT | male | 5 | 21 | 27.1 (24, 31.5) |  | WT+OT vs KO+OT | n.s. |

**Table 6-Supplement 1,2,3.**

| Figure | Parameter | Genotype | Treatment | Gender | Number of individuals | Number of neurons | Mean ± SEM | Statistical test | p-value |
| --- | --- | --- | --- | --- | --- | --- | --- | --- | --- |
| 6supl1 C | Glut frequency (Hz) | WT | - | male | 5 | 13 | 0.73 ± 0.2 | Mann-Whitney | n.s. |
| | | *Magel2* KO | - | male | 5 | 16 | 0.67 ± 0.14 | | |
| 6supl1 E | GABA frequency (Hz) | WT | - | male | 5 | 12 | 0.30 ± 0.08 | Mann-Whitney | n.s. |
| | | *Magel2* KO | - | male | 5 | 12 | 0.26 ± 0.03 | | |
| 6supl1 B | Glut amplitude (pA) | WT | - | male | 5 | 13 | 33.3 ± 3 | Mann-Whitney | **p<0,01** |
| | | *Magel2* KO | - | male | 5 | 14 | 23.8 ± 1.8 | | |
| 6supl1 D | GABA amplitude (pA) | WT | - | male | 5 | 13 | 17.6 ± 2.7 | Mann-Whitney | n.s. |
| | | *Magel2* KO | - | male | 5 | 11 | 13.5 ± 1 | | |

| Figure | Parameter | Genotype | Treatment | Gender | Number of individuals | Number of neurons | Median (Q1,Q3) | Statistical test | p-value |
| --- | --- | --- | --- | --- | --- | --- | --- | --- | --- |
| 6supl2 C | Total length apical dendrites | WT | - | male | 3 | 7 | 3957 (3349,4372) | Mann-Whitney | p=0.89 |
|  |  | *Magel2* KO | - | male | 3 | 11 | 3982 (3472,4760) |  |  |
| 6supl2 D | Total length basal dendrites | WT | - | male | 3 | 7 | 4914 (4102,5739) | Mann-Whitney | p=0.46 |
|  |  | *Magel2* KO | - | male | 3 | 11 | 5596 (4546,6318) |  |  |
| 6supl2 E | Mean number of apical bifurcations | WT | - | male | 3 | 7 | 30 (22,36) | Mann-Whitney | p=0.31 |
|  |  | *Magel2* KO | - | male | 3 | 11 | 32 (31,37) |  |  |
| 6supl2 F | Mean number of basal bifurcations | WT | - | male | 3 | 7 | 35 (30,38) | Mann-Whitney | p=0.25 |
|  |  | *Magel2* KO | - | male | 3 | 11 | 30 (25,33) |  |  |
| 6supl2 G | Mean number of apical branches | WT | - | male | 3 | 7 | 61 (45,74) | Mann-Whitney | p=0.31 |
|  |  | *Magel2* KO | - | male | 3 | 11 | 65 (64,75) |  |  |
| 6supl2 H | Mean number of basal branches | WT | - | male | 3 | 7 | 74 (64,80) | Mann-Whitney | p=0.29 |
|  |  | *Magel2* KO | - | male | 3 | 11 | 64 (70,84) |  |  |

**Table 7.**

| Figure | Parameter | Days in vitro | Genotype | Treatment | Number of preparations | Number of neurons | Mean ± SEM | Statistical test | p-value |
| --- | --- | --- | --- | --- | --- | --- | --- | --- | --- |
| 7A | Percentage of responsive neurons | DIV2 | WT | - | 3 | 169 | 57.88 ± 6.866 | Unpaired t test with Wlech's correction | p=0.479 |
| | | | *Magel2* KO | - | 4 | 210 | 64.15 ± 5.519 | | |
| | | DIV4 | WT | - | 5 | 330 | 29.42 ± 4.315 | | **p<0.0001** |
| | | | *Magel2* KO | - | 3 | 175 | 62.09 ± 5.196 | | |
| | | DIV8 | WT | - | 3 | 206 | 2.553 ± 1.052 | | p=0.233 |
| | | | *Magel2* KO | - | 2 | 179 | 1.016 ± 0.709 | | |
| | | DIV11 | WT | - | 2 | 92 | 1.759 ± 1.188 | | p=0.813 |
| | | | *Magel2* KO | - | 2 | 97 | 1.333 ± 1.333 | | |
| Figure | Parameter | Age | Genotype | Treatment | Number of individuals | Number of neurons | Mean ± SEM | Statistical test | p-value |
| 7C | Driving Force GABA (mV) | P1 | WT | - | 3 | 19 | 12.7 ± 1.5 | Unpaired t test with Wlech's correction | p=0.0535 |
| | | | *Magel2* KO | - | 3 | 20 | 16.8 ± 1.4 | | |
| | | P7 | WT | - | 6 | 42 | 5.0 ± 1.4 | | **p=0.0358** |
| | | | *Magel2* KO | - | 7 | 56 | 8.9 ± 1.2 | | |
| | | P15 | WT | - | 3 | 23 | 2.7 ± 1.6 | | p=0.6805 |
| | | | *Magel2* KO | - | 4 | 29 | 1.9 ± 1.1 | | |
| 7D | Driving Force GABA (mV) | P7 | WT | vehicle | 3 | 37 | 8.5 ± 1.3 | one-way ANOVA, Dunnett's post-hoc test | |
| | | | WT | OT | 3 | 37 | 3.5 ± 1.2 | | **WT-vehicle vs WT+OT, p=0.0041** |
| | | | *Magel2* KO | OT | 4 | 56 | 4.1 ± 0.8 | | **WT-vehicle vs *Magel2* KO+OT, p=0.0055** |

**Table 7-Supplement 1.**
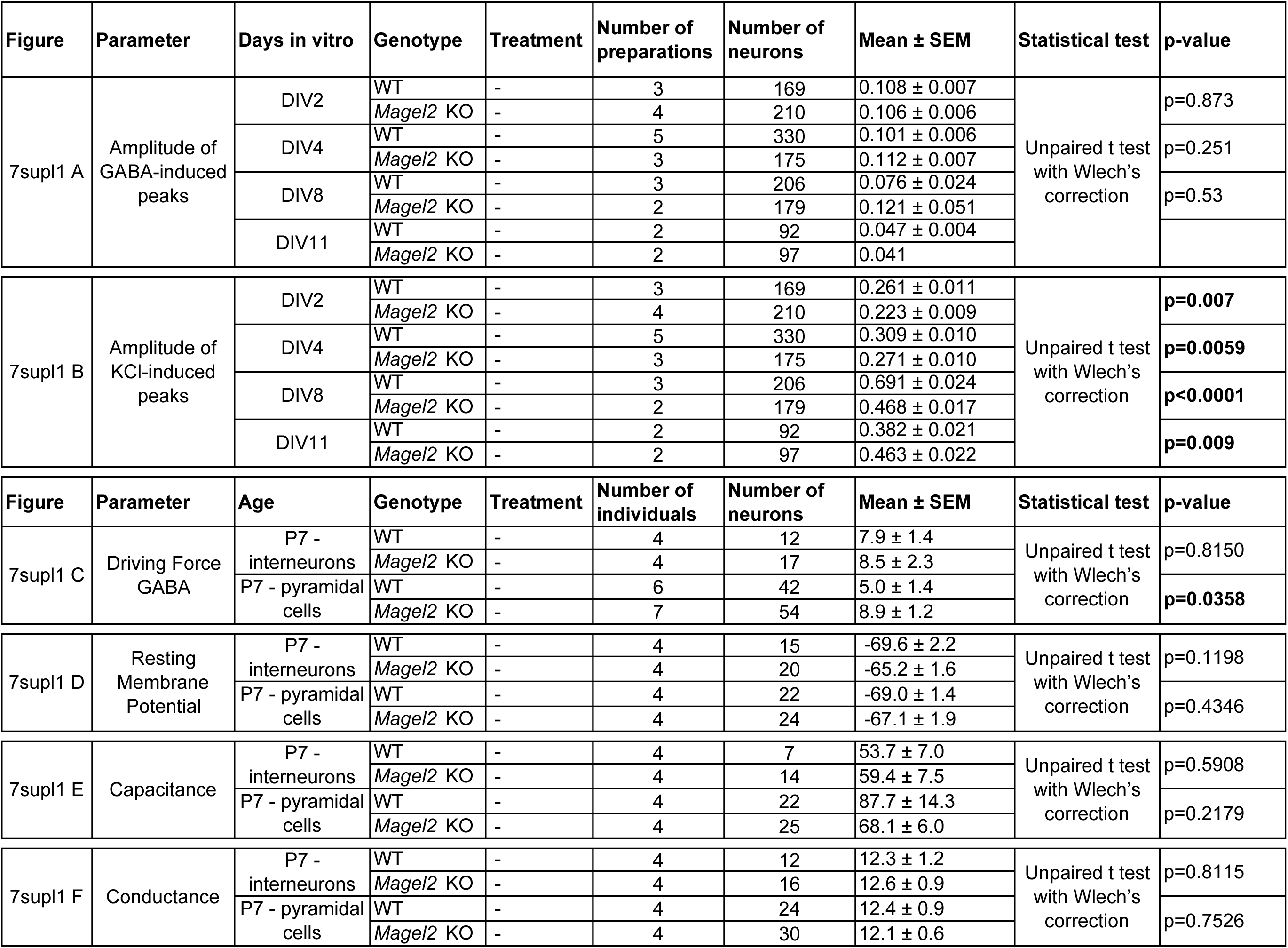

**Table 8.**

| Figure | Parameter | Ratio | Genotype | Treatment | Number of individuals (brains) | Median (Q1, Q3) | Statistical test | p-value |
| --- | --- | --- | --- | --- | --- | --- | --- | --- |
| 8B | Ratio of protein quantity | KCC2/tubulin | WT | - | 5 | 5 (4.8, 5.1) | Mann Whitney | n.s. |
|  |  |  | *Magel2* KO | - | 6 | 4.9(4.5, 5) |  |  |
|  |  | P-Ser<sup>940</sup>/KCC2 | WT | - | 5 | 0.42 (0.21, 0.54) |  | **p<0,05** |
|  |  |  | *Magel2* KO | - | 6 | 0.16 (0.14, 0.20) |  |  |
|  |  | P-Thr<sup>1007</sup>/KCC2 | WT | - | 5 | 0.36 (0.34, 0.66) |  | n.s. |
|  |  |  | *Magel2* KO | - | 6 | 0.49 (0.40, 0.50) |  |  |

